# A stem cell knockout village reveals lineage rewiring and a non-canonical islet cell fate in monogenic diabetes

**DOI:** 10.64898/2025.12.23.696311

**Authors:** Dingyu Liu, Bicna Song, Zhaoheng Li, Stephen Zhang, Tabassum Fabiha, Jiahui Zhao, Ayaka Inoki, Julie Piccand, Chew-Li Soh, Gary Dixon, Aaron Zhong, Nan Hu, Renhe Luo, Batu Ozlusen, Vipin Menon, Ting Zhou, Xiaojie Qiu, Gerard Gradwohl, Dapeng Yang, Kushal Dey, Wei Sun, Wei Li, Danwei Huangfu

## Abstract

Genetics studies have identified a core set of regulators essential for pancreatic β cell development, many of which are mutated in monogenic diabetes. However, how these mutations alter developmental trajectories to produce pathological cell states remains elusive. Here we introduce a knockout village framework that enables longitudinal scRNA-seq profiling of 79 human pluripotent stem cell mutant lines targeting 30 developmental regulators, including 15 diabetes genes, across five islet differentiation stages. We show that loss of lineage regulators impairs β cell formation in a stage-specific manner and rewires developmental trajectories towards competing lineages. Notably, several monogenic diabetes gene mutations drive a shift from β cells to enterochromaffin (EC)-like cells, a recently recognized non-canonical islet cell fate. These EC-like cells exhibit incomplete activation of hormone regulation programs, along with elevated neuron signatures. Leveraging the diversity of cell fate outcomes across mutants, we predicted and experimentally validated ISL1 as a key downstream effector of PDX1 and PAX6 that safeguards β cell identity against an EC-like fate. Together, our findings reveal cell fate rewiring as a widespread, previously underappreciated pathological mechanism in monogenic diabetes and establish a scalable platform for uncovering developmental vulnerabilities in human genetic disorders.

## Introduction

Cell-fate specification during development relies on precisely controlled activity of lineage regulators, and mutations in these factors underlie many developmental disorders [1, 2]. Although single-cell atlases have comprehensively charted their expression dynamics, how genetic mutations in these regulators causally affect developmental trajectories remains largely unexplored. This gap reflects a limitation of current experimental systems, which generally lack the scale and temporal resolution needed to model patient-relevant mutations in a high-throughput manner while tracking their effects on cell-fate transitions over time.

During pancreatic β-cell development, many regulators have been identified that orchestrate successive stages of lineage specifications, from gastrulation to gut tube formation, pancreas budding, and endocrine specification [3–5]. Germline loss-of-function (LOF) mutations in these genes cause monogenic diabetes, including neonatal diabetes and maturity onset diabetes of the young (MODY), primarily through reducing the number or impairing the function of β cells [6, 7]. However, this β-cell-centric model does not explain the existence of pathological cell states with unknown identities (e.g., hormone-deficient endocrine cells) in certain patients and animal models [8, 9]. This raises the broader, unsolved question: does the loss of lineage regulators only block the progression of β cell specification, or can this loss also alter broader differentiation trajectories and contribute to unexplored clinical phenotypes?

Human pluripotent stem cell (hPSC) differentiation systems combined with genome editing provide a powerful platform to model genetic perturbations in human islet development [10, 11]. However, differentiation batch effects, clonal variations, and the labor-intensive nature of profiling individual mutant lines often confound biological interpretations and constrain throughput. While CRISPR-based single-cell screens (e.g., Perturb-seq) [12–14] have expanded the scalability of interrogating disease-relevant genes in hPSC differentiation models, variable gRNA performance can limit precise modeling of patient-relevant genotypes, such as monoallelic or biallelic LOF mutations, thereby constraining direct attribution to patient-specific pathological changes. Recent stem cell ‘village’ strategies, which pool cells from multiple donors for shared differentiation and single-cell profiling, have proven effective in linking genetic variations to cellular phenotypes [15–19]. We reasoned that extending this framework from correlations of natural variation to assessments of defined, engineered genotypes would enable scalable, causal resolution of how specific genetic alterations shape cell fate.

Here we developed a knockout village approach that pools gene-edited isogenic hPSC lines, rather than donor-derived lines, during islet differentiation to dissect the roles of 30 key pancreatic developmental regulators, including 15 diabetes genes. Our knockout village comprised 79 uniquely barcoded, genotype-verified hPSC lines and was profiled by scRNA-seq across five islet differentiation stages. Systematic analysis of cell types, developmental trajectories, and gene programs revealed a broader paradigm for monogenic diabetes in which loss of lineage regulators frequently shifts cells toward developmentally related competing lineages rather than merely arresting cells at earlier progenitor stages. Notably, multiple monogenic diabetes mutants converged at an increased prevalence of enterochromaffin (EC)-like cells, resembling the hormone-deficient cells observed in patients with *RFX6* mutations [8, 9]. By delineating gene programs of this EC-like non-canonical cell fate from closely related canonical islet cell types, we predicted previously unstudied regulators and experimentally validated ISL1, which acts downstream of PDX1 and PAX6 to enforce β cell fate and suppress EC-like identity. These findings demonstrate that diabetes genes safeguards β cell differentiation against sequential fate deviations toward competing lineages, providing new mechanistic insights and avenues for precision medicine.

## Results

### Time-resolved scRNA-seq profiling of a knockout village in human islet differentiation

We selected 30 genes to capture a spectrum of diabetes relevance, from genes causally linked to neonatal diabetes and MODY to genes inferred to influence type 2 diabetes (T2D) risk [6, 20, 21], together with developmental regulators whose LOF mutations lead to disease-relevant phenotypes in mouse pancreas development or hPSC-derived islet differentiation models [22–27] (Fig.1a). These genes are expressed across diverse stages of islet development (Supplementary Fig. S1a), enabling the interrogation of potential differentiation trajectory rewiring over a broad developmental window. To model patient-relevant LOF mutations, we generated 76 Cas9-edited mutant hPSC lines in 2 cell backgrounds (H1 and HUES8) (Supplementary Table.S1). Most clones carried biallelic LOF frameshift mutations, with the remaining harboring monoallelic LOF mutations or enhancer deletions that reduced target gene expression (Supplementary Fig.S1b). To streamline interpretation, clones sharing the same target gene and mutation type were grouped as a single genotype. In total, there were 36 genotypes, 26 of which were represented by multiple clones, allowing assessment of phenotypic reproducibility across clones.

**Fig. 1.**
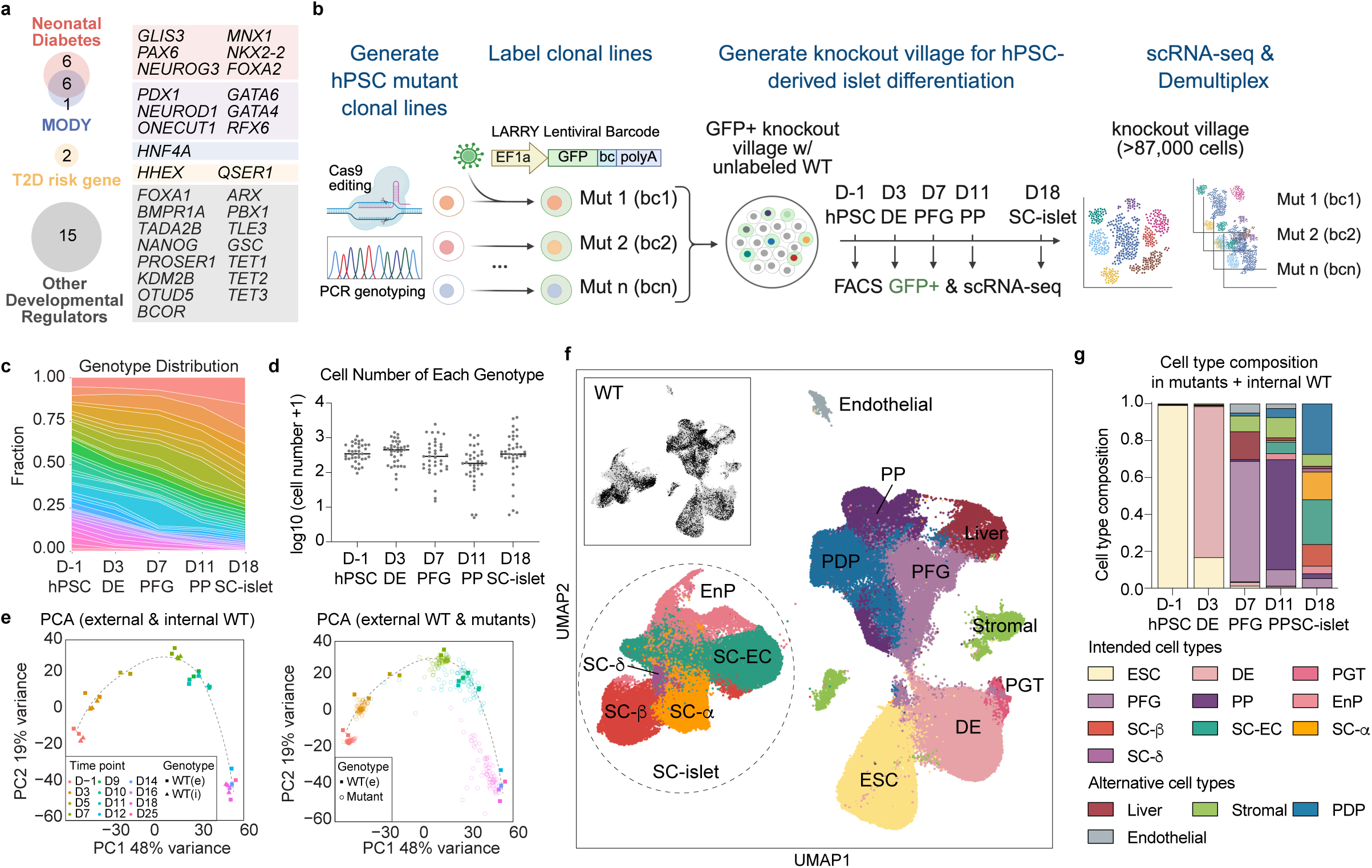
A knockout village of 79 hPSC lines during islet differentiation. **a.** The list of perturbed genes, grouped by the disease relevance. MODY: maturity-onset diabetes of the young. T2D: type 2 diabetes. **b.** Experimental overview of the knockout village. **c.** Stacked line plot showing the genotype representation during differentiation. Each line represents a genotype. **d.** Cell number of individual genotypes during differentiation. The lines represent the median cell number. **e.** Principal component analysis (PCA) of pseudo-bulk scRNA-seq samples from external and internal WT samples (left) and external WT and mutant samples (right). Each dot represents a clone, colored by collection time point. WT(i): internal WT cells pooled and differentiated in the knockout village. WT(e): external WT cells differentiated independently as a reference. **f.** UMAP embedding of scRNA-seq dataset from all time points, annotated by cell types. The distribution of WT cells (both external and internal) is shown on the top. **g.** Cell type composition of the knockout village during differentiation. ESC: embryonic stem cell. DE: definitive endoderm. PGT: primitive gut tube. PFG: posterior foregut. PP: pancreatic progenitor. EnP: endocrine precursor. SC-EC: stem cell-derived enterochromaffin cells. PDP: pancreatic-duodenal progenitor.

To increase scalability and reduce batch variation, we implemented a village strategy. Each hPSC line (76 mutants and three internal wildtype (WT) controls) was uniquely labeled with a constitutively expressed GFP-barcode (cloned from the LARRY library [28]) and then pooled at equal ratios (Fig.1b). During islet differentiation, the pooled village was co-cultured with excess unlabeled WT cells (30% village : 70% WT) to reduce potential non-cell-autonomous interactions among mutants. The village cells were then isolated by fluorescence-activated cell sorting and subjected to scRNA-seq at five key differentiation stages: day -1 (hPSC, 1 day prior differentiation), day 3 (definitive endoderm, DE), day 7 (posterior foregut, PFG), day 11 (pancreatic progenitor, PP) and day 18 (early SC-islet), capturing transitions from pluripotency through gastrulation, gut tube patterning, pancreas budding and endocrine specification. As an independent reference, GFP-labeled WT cells (external WT) were differentiated separately and profiled across a more densely sampled time course.

In total, 87,173 pooled cells passed quality control for both LARRY GFP barcodes and transcriptomes (Supplementary Fig.S1c-d). All clones were captured on day -1, with 201±79 cells each. Nearly all clones were retained throughout differentiation, except for one not detected on day 11 and another on day 18 (Fig.1c-d, Supplementary Fig.S1e-f). Pseudo-bulk analysis showed that internal and external WT clustered closely at all stages (Fig.1e), indicating that the knockout village platform didn’t significantly affect internal WT differentiation. Therefore, we used internal WT as controls for the remaining analysis. In contrast, many mutant clones diverged from the WT trajectory, suggesting differentiation defects. To quantify phenotypic variations, we modeled time- and clone-specific pseudo-bulk gene expression as a function of read depth, cell background, and genotype (Supplementary Fig.S1g). Before differentiation (day -1), when most perturbed genes were not expressed and thus unlikely to contribute, each of these factors explained similar levels of gene expression variance. As differentiation progressed, however, genotype quickly became the dominant contributor, indicating that transcriptomic phenotypes were primarily driven by genetic perturbations.

All expected cell types were observed during islet differentiation, including DE (expressing *EOMES, SOX17*) on day 3, PFG (*ONECUT1, PDX1*) on day 7, PP (*PDX1, SOX9*) on day 11, and SC-islet (*CHGA, NKX2-2*) on day 18, comprising SC-β (*INS, PDX1*), SC-α (*GCG, ARX*), SC-δ (*SST*), SC-enterochromaffin (SC-EC) (*TPH1, SLC18A1*), and their common progenitors, endocrine precursor (EnP) (*NEUROG3*)(Fig.1f-g, Supplementary Fig.S1h-i). In addition, some cells resembled developmentally related alternative lineages, including stromal (*COL3A1*), endothelial (*PLVAP*), liver (*HNF4A, AFP*) and pancreatic-duodenal progenitors (PDP) that co-expressed PP (*HES1, GLIS3*) and duodenal (*CDX2, KLF5*) markers. Together, these results demonstrated that our village platform successfully captured mutant clones across a spectrum of developmental stages and cell types.

### Loss of lineage regulators impairs β cell formation and/or transcriptomic states

Diabetes arises from the loss or dysfunction of insulin-producing β cells. Given that half of the mutated genes are directly linked to diabetes (Fig.1a), we sought to determine the primary, and potentially the earliest, pathological alterations of β cells in each mutant genotype. Clones of the same genotype exhibited highly consistent cell type compositions, suggesting reproducible phenotypes (Supplementary Fig.S2a-c, Supplementary Table.S2). We therefore applied a clone-level linear regression model, accounting for both cell background and target gene, to quantify genotype-specific perturbation effects on cell type composition. We first examined β cell formation by measuring SC-β cell fraction in each genotype with sufficient cell representation (>25 cells on day 18; Fig.2a). Most mutants showed some degree of SC-β depletion, and around 35% (11/32) displayed a significant reduction relative to WT, indicating defects in β cell specification.

**Fig. 2.**
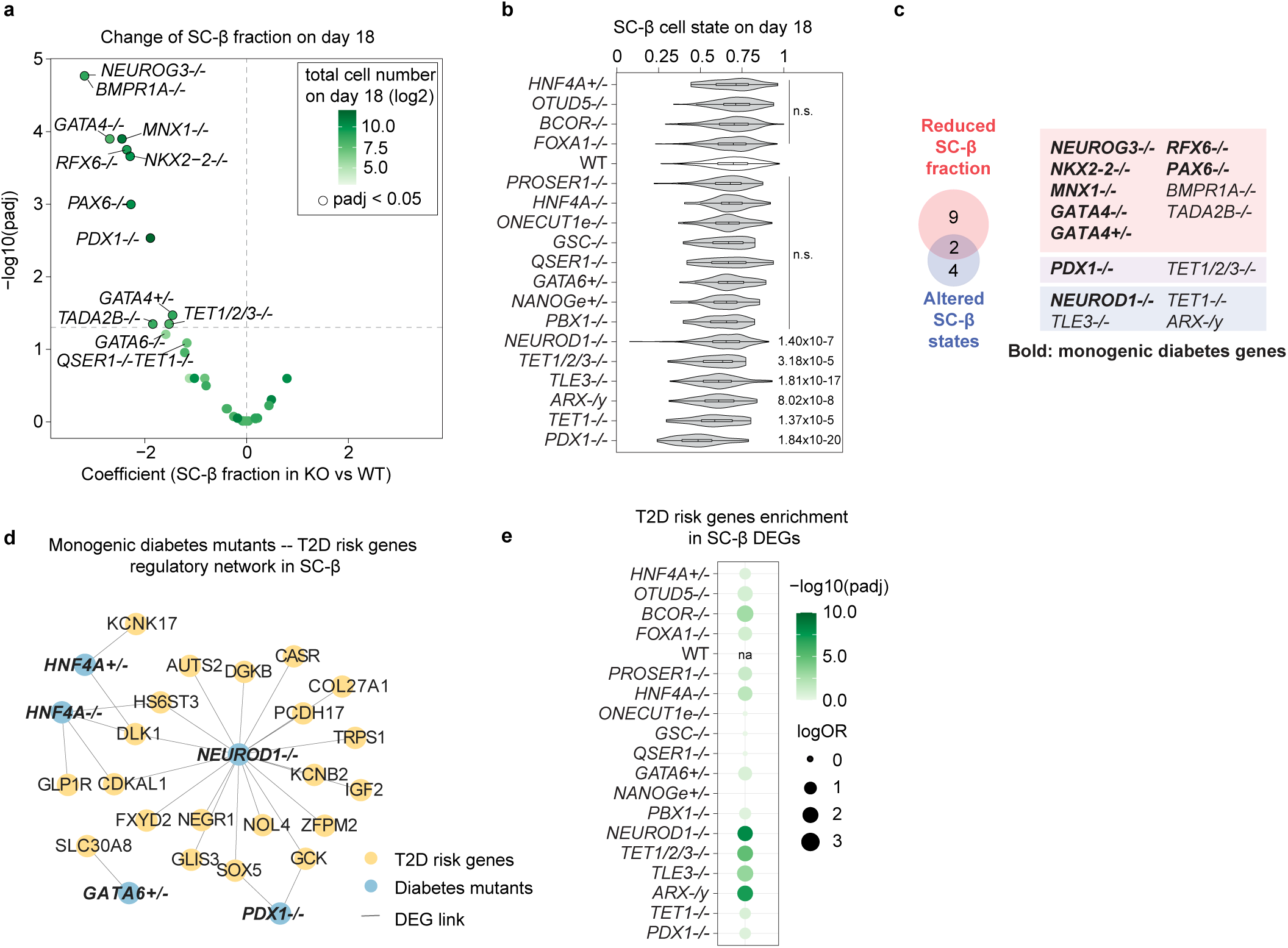
Loss of lineage regulators impairs β cell formation and β cell state. **a.** Volcano plot showing the impact of mutant genotypes on SC-β fraction on day 18. Each dot represents a mutant genotype with over 25 cells on day 18, colored by the total cell number on day 18. **b.** SC-β cell state scores in mutants with over 25 SC-β cells on day 18. *P* values were calculated using Wilcoxon rank-sum test with Benjamini–Hochberg correction. **c.** Venn diagram summarizing mutant genotypes with significant SC-β loss or altered SC-β cell state from (a-b). **d.** Network showing monogenic diabetes mutants (blue) and their differentially expressed genes (DEGs, yellow) that are inferred as T2D risk genes in Open Targets database (Locus-to-Gene score > 0.5). Lines connect mutants to their DEGs. **e.** Enrichment of T2D risk genes among DEGs of individual mutant genotypes in SC-β cells. Enrichment was assessed by Fisher’s exact test with Benjamini–Hochberg correction.

For 18 genotypes that generated an adequate number of SC-β cells per clonal line (>25 SC-β cells), we further assessed potential changes in cell state as a proxy for functional alterations. β cell state was quantified using a similarity score (range between 0-1) derived from our Perturbation-response Score (PS) model [29], computed from the expression of 95 SC-β-specific genes (Supplementary Table.S3). Higher scores indicate greater resemblance to WT SC-β expression patterns, whereas lower scores reflect increasing deviation from the WT state (Fig.2b). Six genotypes, including *PDX1-/-* and *NEUROD1-/-*, showed significantly reduced scores, suggesting SC-β cell state perturbations (Fig.2b, Supplementary Fig.S3a-b). Notably, *NEUROD1-/-* exhibited normal SC-β cell proportion yet significant transcriptomic divergence. This aligned with murine findings that loss of *Neurod1* impaired β cell state and perinatal proliferation without affecting initial specification [30–32]. In support of this, gene set enrichment analysis (GSEA) revealed that transcriptomic alteration in *NEUROD1-/-* SC-β resembled those in *Neurod1-/-* mouse endocrine cells at E15.5 and E17 (Supplementary Fig.S3c) [30, 32]. Ion transport genes crucial for β cell function (e.g., *RYR2, ATP2B4, KCNJ6*) were downregulated, while neuronal sodium transport regulators (e.g., *NKAIN2, NKAIN3*) and lipoprotein genes (e.g., *APOA2, APOB, LIPC*) were aberrantly upregulated (Supplementary Fig. S3a).

Together, these results show that many mutations impaired β cell specification (Fig.2c), while some also disrupted essential β-cell functional programs. A recent integrative study identified monogenic diabetes gene *RFX6* as a regulatory hub in early T2D [33], suggesting the crosstalk between monogenic and polygenic diabetes genes. To explore whether such convergence is already evident during development, we evaluated the enrichment of T2D risk genes among the differentially expressed genes in mutant SC-β cells. We curated 485 T2D-associated genes from Open Targets (Locus-to-Gene score > 0.5; Supplementary Table S4), and they were significantly enriched in differentially expressed genes of mutant lines (odds ratio = 5.11, *p* = 5.47 × 10^-14^), suggesting that monogenic diabetes genes and developmental regulators already influence T2D-relevant network during β cell differentiation (Fig. 2d). *NEUROD1-/-, TET1/2/3-/-, ARX-/y*, and *TLE3-/-* showed the strongest enrichment (Fig. 2e), underscoring their central roles in establishing T2D-related regulatory architecture. In summary, our findings reveal a convergence of monogenic and polygenic diabetes genes at the level of gene regulatory network (GRN) in differentiating β cells, emphasizing that early developmental disruptions can shape adult disease risk.

### Temporal mapping uncovers stage-specific islet differentiation defects and widespread lineage rewiring

To pinpoint when developmental defects first arise, we assessed the differentiation efficiency of mutant genotypes towards all intended cell types along the β cell trajectory. We identified that mutants disrupted β cell differentiation in a stage-specific manner (Fig.3a). Two genotypes (*GATA6-/-* and *GSC-/-*) showed significantly decreased DE cells as early as day 3, indicating gastrulation defects. Nine genotypes (e.g. *FOXA2-/-, HHEX-/-, GATA4-/-*) displayed reduced PFG cells on day 7, indicating impaired gut tube formation and pancreas budding. Five (e.g. *NEUROG3-/-, RFX6-/-, NKX2-2-/-*) showed reduced SC-islet cells, underscoring the requirement of these genes for endocrine specification. Four genotypes (*MNX1-/-, PAX6-/-, PDX1-/-*, *TADA2B-/-*) specifically impaired SC-β cells, indicating these genes are essential for β cell formation. In most cases, early-stage impairments propagated through development, ultimately resulting in reduced SC-islet population on day 18. In extreme cases (e.g., *HHEX-/-*, *HHEX+/-*), mutant genotypes caused a severe growth disadvantage, leading to pronounced cell dropout during subsequent stages of differentiation.

**Fig. 3.**
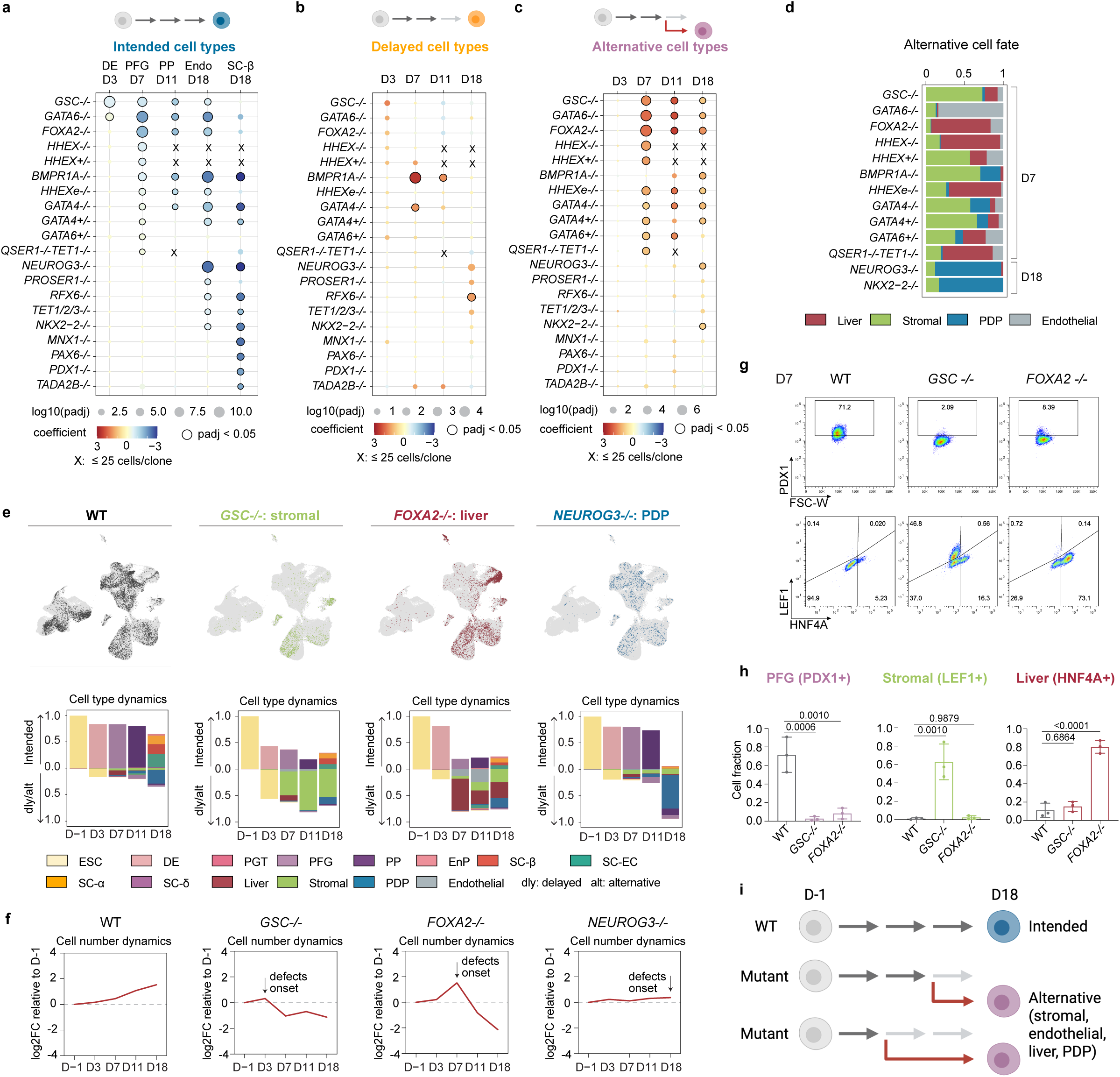
Impaired islet differentiation is accompanied by increase of alternative lineages. **a.** Dot plot showing the impact of mutant genotypes on intended cell types. Endo represents all endocrine cells on day 18, including SC-β, SC-α, SC-δ, SC-EC and EnP. Each mutant genotype was compared to WT from the corresponding time point and cell type following a linear regression model. Genotypes with significant cell type composition changes (p_adj<0.05) are marked as outlined dots. Genotypes for which all clones contained 25 or fewer cells are marked with an ‘x’ (not analyzed). **b.** Dot plot showing the impact of mutant genotypes on delayed cell types. Delayed cells refer to intended cell types captured at later-than-expected time points (e.g., ESC and DE cells on day 7). **c.** Dot plot showing the impact of mutant genotypes on alternative cell types. Alternative cell types refer to non-endoderm (stromal, endothelial), non-pancreatic (liver-like), and non-endocrine (PDP) cell. **d.** Stacked bar graph showing the fraction of individual alternative cell types among all alternative cells in mutant genotypes that exhibited a significant increase in alternative cell types in (c). The time point at which the quantification was performed is indicated on the side. **e.** UMAP (top) and stacked bar plot (bottom) of WT and representative mutant genotypes showing biases towards stromal (*GSC-/-*), liver (*FOXA2-/-*) and PDP (*NEUROG3-/-*) lineages. Dly: delayed differentiation. Alt: alternative lineages. **f.** The dynamics of total cell numbers relative to D-1 in WT, *GSC-/-, FOXA2-/-,* and *NEUROG3-/-* lines. **g.** Representative flow cytometry plots for PFG (PDX1), stromal (LEF1) and liver-like (HNF4A) markers in WT, *GSC-/-*, and *FOXA2-/-* cells on day 7. **h.** Quantification of flow cytometry analysis. n = 3 independent differentiations. *P* values were calculated using one-way ANOVA test with Dunnett’s multiple comparisons test. **i.** Schematic illustrating that loss of lineage regulators disrupts intended islet differentiation in a stage-specific manner and divers progenitors into developmentally related alternative lineages, including stromal, endothelial, liver, and PDP fates.

In addition to revealing differentiation defects in the intended islet lineage, the longitudinal scRNA-seq data also enabled systematic mapping of mutants’ ultimate fate decisions. Surprisingly, only three mutant genotypes displayed a significant delay in differentiation, marked by persistence of earlier intended cell types at later time points (e.g., DE cells on day 11) (Fig.3b). In contrast, most mutant genotypes showed significant increases of alternative cell fates from day 7 onward, suggesting lineage rewiring rather than mere developmental delay (Fig.3c). Moreover, these alternative cell fates are not random but correspond to branch points in islet differentiation, including stromal/endothelial (non-endoderm), liver (non-pancreas), and PDP (non-endocrine) cells (Fig.3d). Loss of different regulators shifted cells towards distinct competing fates. For example, *GSC-/-* led to accumulation of hPSCs on day 3, which subsequently adopted stromal fates, likely reflecting a failure of endoderm commitment (Fig.3e). *FOXA2-/-* caused increased liver cells from day 7 onward, suggesting its requirements in pancreas specifications. *NEUROG3-/-* resulted in depletion of endocrine cells and increase of non-endocrine PDP cells on day 18. Notably, mutant cell representation was not markedly reduced at defect onset (Fig.3f), suggesting that the increased fraction of alternative cell types reflects active fate rewiring rather than a passive consequence of depletion of the intended cell types. In support of scRNA-seq results, individual differentiations confirmed that both *GSC-/-* and *FOXA2-/-* produced fewer PDX1+ PFG cells on day 7 (Fig.3g-h). *GSC-/-* primarily adopted LEF1+ stromal fate, whereas *FOXA2-/-* favored HNF4A+ liver fate. Collectively, these results demonstrate that loss of lineage regulators not only disrupts islet specification but also actively rewires developmental trajectories (Fig.3i), underscoring their central roles in enforcing lineage commitment and constraining cellular plasticity during development.

### PDX1, RFX6, and PAX6 suppress SC-EC fate in islet differentiation

Despite the profound loss of SC-β cells, some mutant genotypes (e.g., *PDX1-/-, PAX6-/-, MNX1-/-*) retained a total endocrine fraction comparable to WT (Fig.4a), suggesting skewed cell type composition within SC-islets. Indeed, *MNX1-/-* resulted in an increase of SC-δ cells (Supplementary Fig.S4a), consistent with its role in mouse islet development [34]. *NKX2-2-/-* showed a bias towards EnP and SC-α cells, concordant with murine finding that *Nkx2-2-/-* endocrine cells were arrested at an incompletely differentiated state [35]. Surprisingly, multiple mutant genotypes, including *PDX1-/-, RFX6-/-*, and *PAX6-/-*, led to a marked increase of SC-EC cells at the expense of SC-β cells (Fig.4b-d). SC-EC markers (*SLC18A1, TPH1, LMX1A*) were upregulated, accompanied by broad downregulation of pancreatic hormone genes (Fig.4c). To confirm the scRNA-seq observations, we examined endocrine cell type compositions in individually differentiated mutant lines on day 18 (Fig.4e, Supplementary Fig.S5a). Consistent with the scRNA-results (Fig.4a), the total CHGA+ endocrine fraction was significantly reduced in *RFX6-/-* but not in *PDX1-/-* or *PAX6-/-* (Fig.4e, Supplementary Fig.S5b). C-PEP+NKX6-1+ SC-β cells, GCG+ SC-α cells and SST+ SC-δ cells were significantly depleted across all genotypes, while SLC18A1+ SC-EC cells were greatly expanded in *RFX6-/-* and *PDX1-/-,* with a similar upward trend in *PAX6-/-* (Fig.4f). When assessed cell fate bias within CHGA+ endocrine cells, all mutants showed a significant increase in SLC18A1+/CHGA+ ratio, with the number reaching around one in *RFX6-/-* and *PDX1-/-*, indicating nearly all endocrine cells became SC-EC cells (Fig.4f-g). Together, our results reveal a previously unrecognized role of PDX1, RFX6, and PAX6 in suppressing SC-EC cells during islet development.

**Fig. 4.**
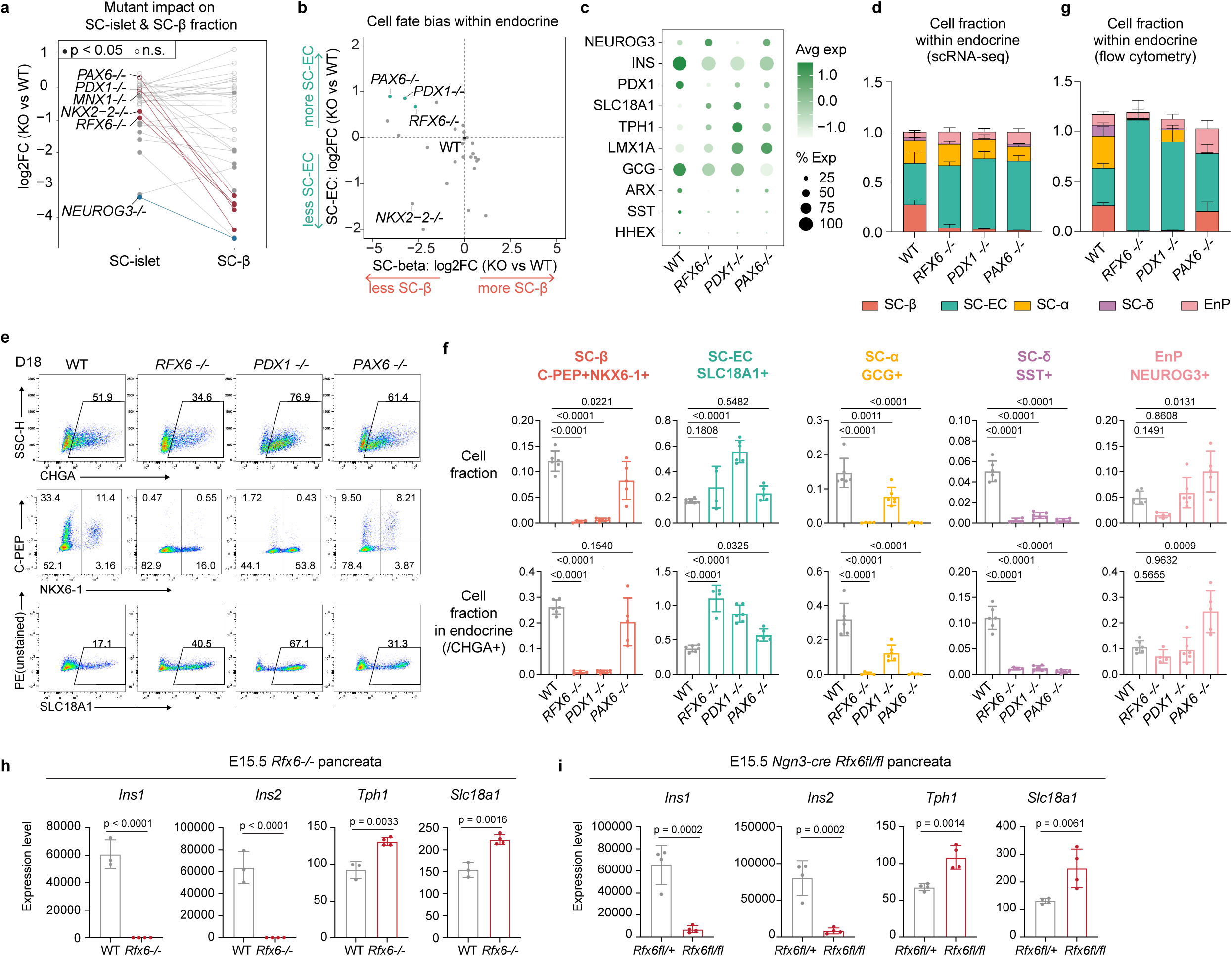
Loss of *RFX6*, *PDX1,* or *PAX6* increases SC-EC cells at the expense of SC-β cells on day 18. **a.** Log2 fold change of SC-islet and SC-β cell fractions on day 18 in mutant genotypes relative to WT. SC-islet represents the entire endocrine population, including SC-β, SC-α, SC-δ, SC-EC and EnP. Each dot represents a mutant genotype; filled dot indicates a significant change (p < 0.05, from cell type composition regression test). **b.** Scatter plot summarizing log2 fold change of SC-β and SC-EC cell fractions within endocrine population in mutant genotypes relative to WT. **c.** Expression of endocrine cell type markers in endocrine cells from WT and mutant genotypes on day 18. **d.** Cell type composition (scRNA-seq) within endocrine population for WT, *RFX6-/-*, *PDX1-/-*, and *PAX6-/-* on day 18. Error bars represent the standard deviation across clones from the same genotype. **e.** Representative flow cytometry plots for pan-endocrine (CHGA), SC-β (C-PEP, NKX6-1) and SC-EC (SLC18A1) markers in WT, *RFX6-/-*, *PDX1-/-* and *PAX6-/-* cells on day 18. **f.** Quantification of flow cytometry analysis. Top: cell type composition on day 18. Bottom: cell type composition within total endocrine cells (CHGA+) on day 18. n = 6 (WT and *PDX1-/-*), n=5 (*PAX6-/-*) and n=4 (*RFX6-/-*) independent differentiations. *p* values were calculated using one-way ANOVA test with Dunnett’s multiple comparisons test. **g.** Cell type composition (flow cytometry) within endocrine population for WT, *RFX6-/-*, *PDX1-/-*, and *PAX6-/-* on day 18. Error bars represent the standard deviation across differentiation replicates. **h-i.** mRNA level of β cell (*Ins1*, *Ins2*) and EC cell (*Tph1*, *Slc18a1*) markers in E15.5 WT and *Rfx6-/-* pancreata (h), and in E15.5 *Ngn3-cre Rfx6fl/+* and *Rfx6fl/fl* pancreata (i), measured by microarray. n = 4 (*Rfx6-/-, Ngn3-cre Rfx6fl/+,* and *Ngn3-cre Rfx6fl/fl*), n=3 (WT) samples.

Since their first discovery in 2019 [36], SC-EC cells have been frequently observed in hPSC-derived islet differentiation studies, yet they are not detected in healthy adult human islets, and their exact identity remains debated. They were initially hypothesized as ‘off-target’ enterochromaffin cells due to the expression of serotonin (5-HT) biosynthesis and secretion genes (e.g., *TPH1, SLC18A1*) [36], but the lack of canonical intestinal markers argues against a classic intestinal lineage. More recently, rare 5-HT+INS+ cells were detected in human fetal, neonatal, and infant pancreata [37], suggesting that they represent a transient endocrine state in developing islets rather than an *in vitro* artifact. Our findings show that although EC-like cells are rare in normal islet development, they can emerge under pathological conditions such as monogenic diabetes. Because monogenic diabetes patient samples are extremely rare, we performed microarray on *Rfx6* knockout murine fetal pancreata at E15.5. Pan-endocrine marker (*Chga*) and all hormone genes (*Ins1, Ins2, Gcg,* and *Sst*) were greatly diminished In *Rfx6-/-* (Fig.4f, Supplementary Fig.S5c). The SC-EC markers (*Tph1*, *Slc18a1*) were detectable in WT pancreata and significantly upregulated in *Rfx6-/-*. A similar increase of SC-EC markers was also observed in a conditional mouse knockout model, *Ngn3-cre Rfx6fl/fl* (Fig.4i, Supplementary Fig.S5d). These results support the presence of EC-like cells in *Rfx6* knockout fetal pancreata. Notably, it was reported that pancreatic tissue from *RFX6*-mutated neonatal diabetes patients had CHGA+ cells lacking insulin, glucagon or somatostatin [8, 9]. Our differentiation results suggest that these hormone-deficient endocrine cells in monogenic diabetes patients correspond to EC-like cells.

### SC-EC cells exhibit incomplete activation of hormone regulation programs and enhanced neuron signatures

The unexpected role of PDX1, RFX6, and PAX6 in suppressing SC-EC cells prompted us to investigate the identity and underlying gene regulation differences between SC-EC cells and canonical islet cell types. To this end, we defined 36 co-varying gene programs during differentiation using the non-negative matrix factorization (NMF) method [38] (Supplementary Fig.S6a, Supplementary Table.S5). Most programs were active in WT and aligned with cell-type features (Supplementary Fig.S6b-d, Supplementary Table.S6). For example, hPSC was characterized by pluripotency, cell cycle and stress response programs. As differentiation progressed, progenitors (DE, PFG, PP) turned on programs associated with extracellular matrix (ECM), morphogenesis, and digestive tract development. Endocrine cells subsequently activated programs related to hormone regulation and cell projection. Alternative lineages, including stromal-, endothelial-, liver-like and PDP cells, displayed distinct activation of muscle contraction, angiogenesis, lipoprotein metabolism and digestive tract development programs, respectively (Supplementary Fig.S6c-d).

We identified nine gene programs engaged in endocrine commitment (Fig.5a-b, Supplementary Fig.S7a). Three programs (h1, h2, i3) were active across all endocrine cell types and enriched for genes in endocrine pancreas development and hormone regulation. The remaining programs showed greater cell-type specificity. The SC-β (i5) program was uniquely enriched for insulin processing (Fig.5c). Programs of canonical islet cell types (SC-β (i5), SC-δ (i8), and SC-α (i1, i2)) were all strongly associated with cellular secretion and hormone regulation, highlighting shared functions. In contrast, the SC-EC (h3) program was more linked to the neuronal system and synaptic pathways. Neuron markers (dopa decarboxylase *DDC*, glutamate receptor subunit *GRIA2*) were activated in SC-EC cells, whereas hormone regulation genes (*INS, ERO1B*) were lowly expressed (Supplementary Fig.S7b). To assess *in vivo* relevance, we mapped the activity of our gene programs in a human fetal single-cell atlas [39] (Fig.5d). Programs of canonical islet cell types were restricted to fetal pancreatic endocrine cells, confirming that our differentiation faithfully recapitulates embryonic islet development. The SC-EC (h3) program, however, was active in both fetal pancreatic endocrine cells and spinal cord neurons, indicating a neuron-like transcriptional signature. Together, these results demonstrate that canonical islet cells share core endocrine programs, while SC-EC cells diverge toward alternative states by activating a neuron-associated gene program.

**Fig. 5.**
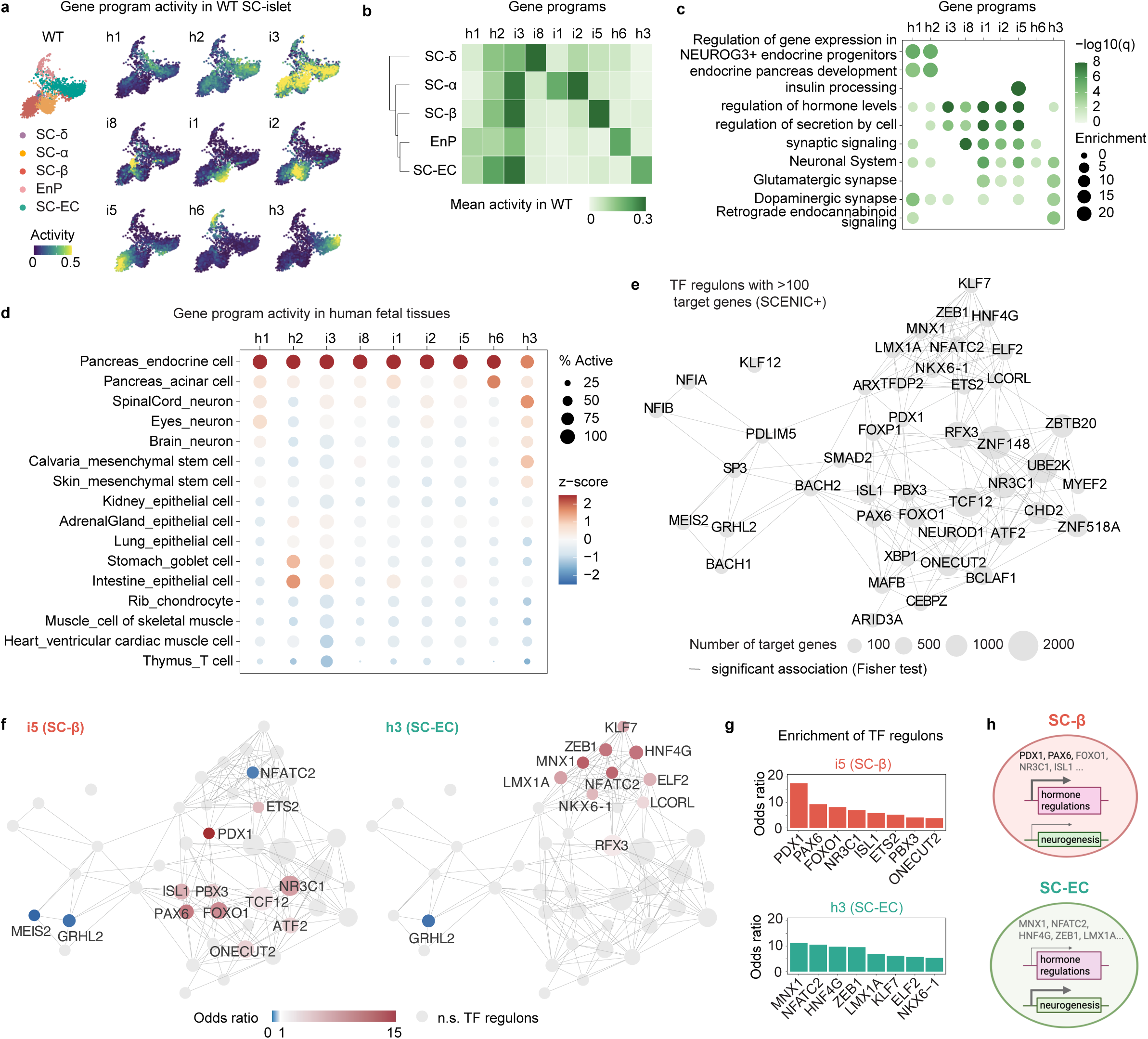
SC-EC cells show reduced hormone regulation program activity and enhanced neuron signatures compared to canonical islet cells. **a.** UMAP embedding of WT SC-islet cells, colored by activity of endocrine-associated gene programs. Gene programs were derived by NMF method on the full scRNA-seq data. **b.** Mean activity of endocrine-associated gene programs across WT endocrine cell types. **c.** Representative gene ontology terms enriched in the top 100 genes of endocrine-associated gene programs. **d.** Relative activity of endocrine-associated gene programs across various cell types in human fetal scRNA-seq data [39]. **e.** Network view of core TF regulons (>100 target genes) inferred by SCENIC+ using WT SC-islet sc-multiome data. Each node represents a TF regulon. Edges connect TF regulons with significantly overlapping target genes. Enrichment was assessed by Fisher’s exact test with Benjamini–Hochberg correction. **f.** Core TF regulons significantly enriched in the top 100 genes of SC-β (i5) and SC-EC (h3) gene programs (*p*<0.001). Enrichment was assessed by Fisher’s exact test with Benjamini–Hochberg correction. **g.** Top eight core TF regulons significantly enriched in SC-β (i5) and SC-EC (h3) gene programs, ranked by odds ratio. **h.** Schematics showing differences between SC-β and SC-EC cell transcriptome states and their candidate upstream transcription factors inferred by SCENIC+.

To identify upstream regulators of SC-β and SC-EC cell states, we inferred transcription factor (TF) regulons using SCENIC+ on a published SC-islet sc-multiome dataset [40, 41]. This analysis discovered 45 core TFs, including known endocrine regulators such as PDX1, PAX6 and ARX (Fig.5e, Supplementary Table.S7). The SC-β (i5) program was strongly enriched for the PDX1 regulon, followed by PAX6, FOXO1, NR3C1 and ISL1 (Fig.5f-g). In contrast, the SC-EC (h3) program was enriched for targets of TFs essential for neuron development, including MNX1, ZEB1, and LMX1A [42–45]. This analysis suggests that SC-β and SC-EC cells are governed by distinct TF regulons (Fig.5h).

### Predictive modeling identified ISL1 as a key repressor of SC-EC cells

While current methods (e.g., SCENIC+) are powerful for predicting TF regulons using WT samples, it remains unclear whether the inferred connections are causal [46–48]. We reasoned that our large cohort of mutant lines could provide an effective reference to refine the causal inference. As a proof-of-concept, we tested whether our data could help prioritize candidate regulators of SC-β and SC-EC programs previously inferred by SCENIC+ (Fig.5f-g). Because both cell types arise from EnP, we hypothesized that the expression of a bona fide regulator in EnP should correlate with the SC-EC/SC-β ratio across mutant genotypes (Fig.6a). To test that, we performed linear regression of all EnP-expressed genes against the SC-EC/SC-β ratio (Fig.6b-c, Supplementary Table.S8). As expected, the SC-EC marker SLC18A1 strongly predicted higher SC-EC/SC-β ratio, while PDX1 and RFX6 showed negative correlations. Among all SCENIC+-inferred core TFs, ISL1 showed the strongest negative correlation with the SC-EC/SC-β ratio (Fig.6b, 6d), suggesting that ISL1 promotes SC-β and suppresses SC-EC differentiation. Notably, ISL1 mutations have not been linked to monogenic diabetes, but have been reported in multiple familial early-onset type 2 diabetes cases [49, 50], further supporting its functional relevance.

**Fig. 6.**
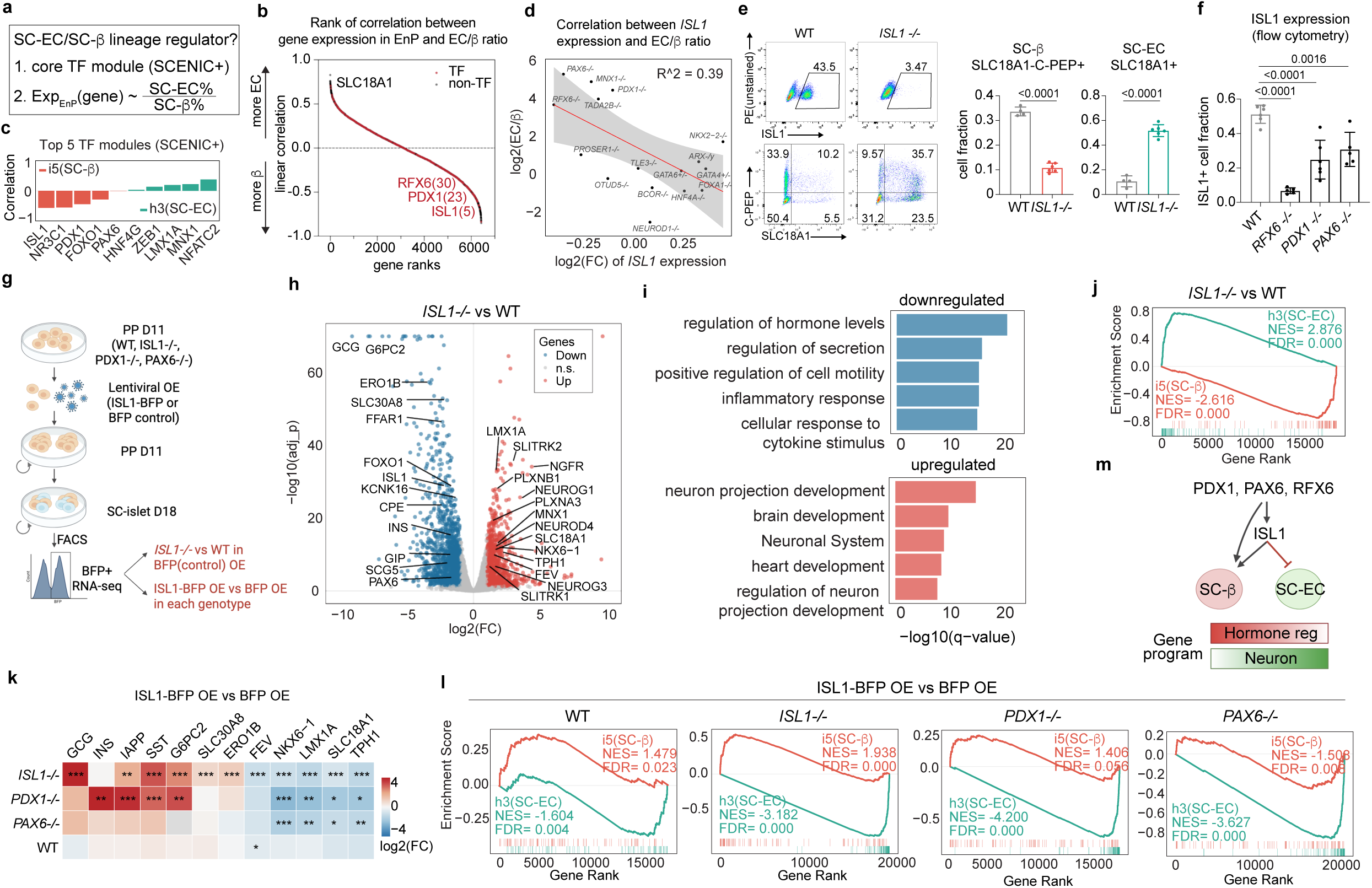
Mutant-based predictive analyses identify ISL1 as a key repressor of SC-EC cells. **a.** Framework for prioritizing the search of candidate regulators of SC-β and SC-EC differentiation. **b.** Waterfall plot summarizing the linear regression coefficients of all EnP-expressed genes. The correlation was computed between the log2 fold change of gene expression in EnP (mutant vs WT) and the SC-EC/SC-β ratio on day 18. **c.** Linear regression coefficients of the top five candidate regulators for the SC-β (i5) and SC-EC (h3) gene programs inferred by SCENIC+. **d.** Linear regression of *ISL1* expression in EnP versus SC-EC/SC-β ratio across mutant genotypes. The solid red line and gray area indicate the regression line and 95% confidence interval, respectively. **e.** Left: representative flow cytometry plots for ISL1, SC-β (C-PEP) and SC-EC (SLC18A1) markers in WT and *ISL1-/-* cells on day 18. Right: quantification of flow cytometry results. n = 6 (*ISL1-/-*), n=4 (WT) independent differentiations. *P* values were calculated using unpaired Welch’s t test. **f.** Quantification of ISL1+ fraction on day 18 from flow cytometry analysis. n = 6 (WT and *PDX1-/-*), n=5 (*RFX6-/-* and *PAX6-/-*) independent differentiations. *P* values were calculated using one-way ANOVA test with Dunnett’s multiple comparisons test. **g.** Schematic of *ISL1* overexpression experiment. **h.** Volcano plot showing differential gene expression in *ISL1-/-* vs WT on day 18. Significance thresholds: abs(log2(FC)) > 1 and p_adjusted < 0.05. **i.** Gene ontology analysis of top five enriched terms among upregulated and downregulated genes in *ISL1-/-* vs WT. **j.** GSEA of *ISL1-/-* vs WT, with genes ranked by log2 fold change and tested for enrichment of top 100 genes of SC-β (i5) and SC-EC (h3) gene programs. **k.** Expression of endocrine cell markers in *ISL1-BFP* OE vs *BFP* OE (control) across different genotype (WT, *ISL1-/-*, *PDX1-/-*, *PAX6-/-*). *** p_adj < 0.001, ** p_adj < 0.01, * p_adj < 0.05. **l.** GSEA of *ISL1-BFP* OE vs *BFP* OE (control) across genotypes, with genes ranked by log2 fold change and tested for enrichment of top 100 genes of SC-β (i5) and SC-EC (h3) gene programs. **m.** Schematic showing the roles of PDX1, RFX6, PAX6, and ISL1 in directing SC-β and SC-EC differentiation.

To validate the predicted role of ISL1, we generated *ISL1-/-* hPSC lines and analyzed cell type compositions on day 18 (Fig.6e). We co-stained C-PEP and SLC18A1 to distinguish SC-β (C-PEP+SLC18A1-) and SC-EC (SLC18A1+) cells. As predicted, *ISL1-/-* resulted in pronounced loss of SC-β cells and gain of SC-EC cells, indicating that ISL1 is essential for safeguarding SC-β specification from SC-EC fate. Moreover, ISL1+ cells were significantly reduced in *PDX1-/-*, *PAX6-/-,* and *RFX6-/-* on day 18 (Fig.6f), suggesting that ISL1 acts downstream of these regulators in endocrine specifications.

To further investigate ISL1 function and its hierarchical relationship with other regulators, we performed ISL1 overexpression (OE) rescue experiments (Fig.6g). *ISL1* (*ISL1-IRES2-BFP* for sorting purpose) was introduced into WT, *ISL1-/-, PDX1-/-*, and *PAX6-/-* cells at the PP stage (day 11) through lentiviral infection, and cells were analyzed on day 18 by bulk RNA-seq. First, compared to WT, *ISL1-/-* showed downregulation of SC-β regulators (*PAX6, FOXO1, ISL1*), hormone genes (*INS, GCG*), secretion machinery (*SLC30A8, G6PC2, ERO1B, SCG5*), and upregulation of SC-EC markers and candidate regulators (*SLC18A1, TPH1, LMX1A*) (Fig.6h). GO analysis confirmed downregulation of hormone secretion pathways and upregulation of neuronal projection programs (Fig.6i), and GSEA further revealed the repression of SC-β (i5) program and the activation of SC-EC (h3) program (Fig.6j). Together, these results established ISL1 as a novel repressor of SC-EC fate in human islet differentiation.

Next, when compared to the *BFP* control, *ISL1* OE repressed neuron projection programs, SC-EC markers and candidate regulators across all mutant genotypes, with a similar downward trend in WT (Fig.6k, Supplementary Fig.S8a-b). Conversely, endocrine genes and programs (*GCG, INS, SST, IAPP* and *G6PC2*) were upregulated, especially in *PDX1-/-* and *ISL1-/-*. The SC-EC (h3) program was significantly repressed across all genotypes, indicating that ISL1 is sufficient to suppress SC-EC cells (Fig.6l). However, the SC-β (i5) program was only robustly induced in *ISL1-/-* and WT but not in *PDX1-/-* or *PAX6-/-*, indicating that PDX1 and PAX6 cooperate with ISL1 to establish SC-β identity. Collectively, our results identified ISL1 as a key downstream effector of RFX6, PDX1, and PAX6 that promotes SC-β fate while constraining SC-EC fate during human islet specification (Fig. 6m). While ISL1 is sufficient to repress the SC-EC fate, full establishment of SC-β programs requires the coordinated action of ISL1 together with PDX1, RFX6, and PAX6.

## Discussion

Understanding how disease genes disrupt human development requires causal, time-resolved mapping of cell fate decisions. Here we present a scRNA-seq village framework that allows longitudinal profiling of cell differentiation in many disease-relevant mutant lines, revealing biologically and clinically meaningful cell states and gene regulatory architectures. We interrogated 30 genes across 79 hPSC lines at five islet differentiation stages in a single experiment. With ongoing advances in gene editing, stem cell and organoid technology, cell culture automation, and resources such as the null allele hPSC collection from the MorPhiC (Molecular Phenotypes of Null Alleles in Cells) Consortium [51], this framework is readily scalable to diverse development and disease contexts. Moreover, such causal, time-resolved data are crucial for training next-generation predictive models of cell state transitions. This is particularly important given the genetic complexity of common diseases, which involve numerous genes and intricate combinatorial effects that preclude exhaustive experimental testing. Current computational models largely rely on descriptive datasets from unperturbed samples [41, 52, 53] and thus far, show poor consistency across methods and struggle to capture causal relationships [46–48]. We demonstrate that integrating mutant-response data with existing prediction methods, even via simple linear regression, can effectively prioritize candidate regulators for validation. Emerging deep learning approaches are now beginning to exploit large-scale perturbation datasets to predict previously unseen cellular responses [54–56]. As these methods rapidly evolve, we expect that causal, time-resolved datasets like ours will improve model accuracy and support the development of virtual cell frameworks [57] for mapping gene function in human biology and disease.

We consistently observed a loss of insulin-producing β cells accompanied by an increase of serotonin-producing EC-like cells across multiple diabetes mutant genotypes. Notably, serotonin production has also been reported in β cells during pregnancy [58] and in pancreatic neuroendocrine tumors [59, 60]. We speculate that this recurrent shift towards a serotonin-producing EC-like state across different compromised conditions is not coincidental but rather reflects an intrinsic vulnerability of the β cell GRN. Gene program analyses suggest that EC-like cells, compared to β cells, exhibit enhanced neuron features. Although β cells and neurons arise from distinct developmental lineages, they share striking similarities in physiology, developmental regulators, and gene expression [61–64]. This relationship may be rooted in their evolutionary history: insulin-like peptides were initially secreted by neurons to regulate metabolism, as seen in organisms such as *Caenorhabditis elegans* and *Drosophila melanogaster*, long before organized islet structures evolved in jawless fish [62, 65]. The propensity of β cells to shift to a more neuron-like EC state suggests that β cells actively maintain a delicate balance between neuron and endocrine programs. We showed that this balance requires sustained engagement of lineage regulators including PDX1, RFX6, PAX6, and ISL1. Previous epigenomic profiling and β-cell-specific mouse knockout studies suggest that epigenetic regulation, particularly PRC2-mediated H3K27me3, also contributes to a neuron-endocrine balance [64, 66]. Future studies are needed to define how transcriptional and epigenetic regulators cooperate to establish the β cell GRN and maintain the neuro-endocrine balance during islet development and homeostasis.

Lineage plasticity is increasingly recognized in cancer and tissue regeneration [67–69], where terminally differentiated cells can escape their original identities and adopt alternative states. In contrast, developmental disorders are often characterized as hypoplasia or malformations, leaving it unclear whether misdirected cell-fate decisions also contribute to disease. Although individual cases of fate redirection have been reported, such as *HHEX* or *ZNF808* deletions producing liver but not pancreatic cells [25, 70] and *Bcl11b* loss diverting T cell precursors toward NK-like cells [71, 72], the overall prevalence of such plasticity has not been systematically assessed. Our findings show that during islet differentiation, lineage regulator perturbations frequently reduce intended cell types while actively diverting progenitors into alternative fates in a gene- and stage-specific manner, suggesting that lineage rewiring may be more generalizable than previously appreciated. Whether those diverted cells are retained in embryos remains uncertain, as there could be *in vivo* surveillance mechanisms that actively eliminate aberrant intermediates [73–75]. In some cases, however, evidence suggests that misdirected cells do persist *in vivo*. For example, both patients and mice with *RFX6* mutations exhibit hormone-deficient endocrine cells that likely correspond to the EC-like cells we observed [8, 9]. This raises the exciting possibility that EC-like cells could serve as an endogenous reservoir for restoring β cells. Indeed, we show that *ISL1* overexpression in monogenic diabetes mutant lines suppresses SC-EC cells and promotes re-establishment of β cell identity. Future efforts to identify and optimize reprogramming strategies may open new therapeutic avenues for this form of monogenic diabetes, and more broadly, demonstrate how developmental plasticity revealed through systematic perturbations can inform regenerative interventions.

## Methods

### I. Experimental procedures

#### hPSC Culture

Mutant and WT hPSCs were maintained in Essential 8 (E8) medium (Thermo Fisher Scientific, A1517001) on vitronectin (Thermo Fisher Scientific, A14700) pre-coated plates at 37 °C with 5% CO2. For regular maintenance, cells were passaged every 3–4 days at 1:10-1:15 ratio using 0.5 mM EDTA (KD Medical, RGE-3130) in PBS for dissociation. The Rho-associated protein kinase (ROCK) inhibitor Y-27632 (10 μM; Selleck Chemicals, S1049) was added to the E8 medium the first day after passaging or thawing of hPSCs. Cells were regularly confirmed to be mycoplasma-free by the Memorial Sloan Kettering Cancer Center (MSKCC) Antibody & Bioresource Core Facility.

#### Generation of mutant hPSC clonal lines in knockout village

Mutant clones were generated on HUES8 (NIHhESC-10-0043) and H1 (NIHhESC-09-0021) dox-inducible Cas9 hPSC lines. Most mutant clonal lines have been previously reported and are listed in the Supplementary Table.1. In brief, for gene knockout clones, a single gRNA targeting an exon was used to induce frameshift mutations. For enhancer deletion clones, a pair of gRNAs flanking the enhancer region was employed to excise the entire enhancer. crRNAs and tracrRNA were ordered from IDT (Alt-R® CRISPR-Cas9 crRNA, #1072532) and diluted to 15 nM final concentration. RNA molecules were transiently transfected into hPSCs using Lipofectamine RNAiMAX (Thermo, 13778100) following manufacturer’s instructions. Cas9 expression was induced with 2 μg/ml doxycycline one day prior to transfection, the day of transfection, and one day after transfection. 3 days after transfection, hPSCs were dissociated to single cells using TrypLE Select (Thermo Fisher Scientific, 12563029), and 500-1,000 cells were plated into one 100-mm tissue culture dish with 10 ml E8 medium supplemented with 10 μM ROCK inhibitor Y-27632 (Selleck Chemicals, S1049) for colony formation. After 7-10 days of expansion, single colonies were picked into 96-well plate. Genomic DNA was extracted using the QIAGEN Blood & Cell Culture DNA Kit (QIAGEN, 13362) for genotyping. Details of the gRNA sequences, PCR primers, and mutation profiles can be found in the Supplementary Table 1.

#### Barcoding and pooling of hPSC clones

Individual GFP-barcode plasmids were cloned from the LARRY barcode library (Addgene #140024) and then transfected to 293T cells to produce lentivirus. Each mutant hPSC clone was then infected with a unique barcode lentivirus, targeting 20-40% GFP positivity. 3 days after infection, GFP+ cells were sorted by flow cytometry, expanded in E8 medium, and cryopreserved in E8 with 10% DMSO (Santa Cruz Biotechnology sc-358801).

To minimize cell viability loss due to handling and waiting time, mutant clones were pooled in two sequential steps. First, barcoded clones were counted and grouped into 11 pools. Each pool contained 5-13 clones, with equal cell numbers contributed per clone. Each clone was assigned to only one pool. Cells were briefly expanded in the pooled format for a few passages, cryopreserved for stock, and immediately processed to avoid skewed cell representation. Second, the 11 pools were counted and mixed to a single village for differentiation as described below.

#### hPSC-directed islet differentiation

hPSCs were seeded at a density of 2.1 × 10^5^ cells/cm2 on Matrigel-coated (Corning, 356230) plates in E8 medium with 10 μM Y-27632. After 24 hours, cells were washed with PBS and differentiated following previously described protocol with some modifications [76].

From day -1 to day 11, cells were cultured on 12-well plates with daily medium change. On day 11, cells were dissociated with TrypLE and transferred into ultra-low attachment 6-well plates (Corning, 3471) at a density of ∼6 × 10^6^ cell per 6-well in 6 ml S5 medium. The plates were incubated on an orbital shaker at 100 rpm. To support cell survival, 10 μM Y-27632 was added to S5 medium on the transfer day. From day 11 to day 17, the medium was changed daily, and afterwards, it was changed every other day.

The differentiation recipes were as follows.

Day 0-2: S1/2 medium supplemented with 100 ng/ml Activin A (Bon Opus Biosciences, C687-1MG) and 5 μM CHIR99021 (Tocris, 4423) for 1 day. S1/2 medium supplemented with 100 ng/ml Activin A for the next 2 days.

Day 3-4: S1/2 medium supplemented with 50 ng/ml KGF (PeproTech, AF-100-19) and 0.25 mM L-Ascorbic acid (Sigma-Aldrich, A4544).

Day 5-6: S3/4 medium supplemented with 50 ng/ml KGF, 0.25 mM L-Ascorbic acid and 1 μM retinoic acid (MilliporeSigma, R2625).

Day 7-10: S3/4 medium supplemented with 50 ng/ml KGF, 0.1 μM retinoic acid, 200 nM LDN (Stemgent, 04-0019), 0.25 μM SANT-1 (Sigma, S4572), 0.25 mM L-Ascorbic acid and 200 nM TPB (EMD Millipore, 565740).

Day 11-17: S5 medium supplemented with 10 µM ALK5i II (Cayman Chemical Company, 14794-5), 0.1 μM retinoic acid, 0.25 μM SANT-1, 0.25 mM VitC, 1 µM T3 (Sigma-Aldrich, T6397), 10 mg/ml Heparin (Sigma-Aldrich, H3149) and 1 µM γ-Secretase Inhibitor XXI (EMD Millipore, 565790).

Day 18 onwards: ESFM medium

The base differentiation medium formulations were as follows.

S1/2 medium: 500 ml MCDB 131 (Cellgro, 15-100-CV) supplemented with 2 ml 45% glucose (MilliporeSigma, G7528), 0.75 g sodium bicarbonate (MilliporeSigma, S5761), 2.5 g BSA (Proliant, 68700), 5.1 ml GlutaMAX (Invitrogen, 35050079).

S3/4 medium: 500 ml MCDB 131 supplemented with 0.52 ml 45% glucose, 0.877 g sodium bicarbonate, 10 g BSA, 2.5 ml ITS-X (Life Technologies, 51500056), 5.2 ml GlutaMAX.

S5 medium: 500 ml MCDB 131 supplemented with 4 ml 45% glucose, 0.877 g sodium bicarbonate, 10 g BSA, 2.5 ml ITS-X, 5.2 ml GlutaMAX.

ESFM medium: 500 ml MCDB 131 supplemented with 0.52 ml 45% glucose, 10.5 g BSA, 5.2 ml GlutaMAX, 5.2 ml NEAA (Invitrogen, 11140050), 1 mM ZnSO4 (MilliporeSigma, 108883), 523 ml Trace Elements A (Corning, 25-021-CI), 523 ml Trace Elements B (Corning, 25-022-CI), 10 mg/ml Heparin.

#### scRNA-seq of knockout village

For differentiation, 11 mutant pools (76 mutant clones) and 3 WT barcoded clones were counted and pooled together to form a knockout village, with equal cell contributions from each clone. To reduce potential non-cell-autonomous interactions among mutants, the knockout village (76 mutant clones + 3 WT clones) was then co-cultured with excess unlabeled WT cells (30% village : 70% WT) during differentiation. Differentiating cells were then collected and cryopreserved in Bambanker cell freezing medium (Fujifilm, 302-14681) at the following time points: day -1 (one day before differentiation), 3, 7, 11 and 18.

On the day of scRNA-seq library preparation, frozen cells were thawed in 37°C and resuspended in PBS with 2% BSA. GFP+ village cells were then sorted out and loaded on Chromium Controller with a targeted collection of 30,000 cells per reaction following the manufacturer’s instructions (10x Genomics Chromium Single Cell 3′ Reagent Kit v3.1 User Guide). cDNA libraries and targeted barcode libraries were generated separately using 10ul cDNA each. cDNA libraries were made following manufacturer’s instructions and targeted LARRY barcode libraries were amplified using specific primers (F: CTACACGACGCTCTTCCGATCT; R: GTGACTGGAGTTCAGACGTGTGCTCTTCCGATCTtaaccgttgctaggagagaccataT). Both cDNA and targeted libraries were sequenced on NovaSeq 6000 platform, with a targeted depth of 20,000 reads/cell for cDNA libraries and 1000 reads/cell for targeted libraries.

#### scRNA-seq of external WT cells

We differentiated and sequenced external WT cells at densely sampled time points to serve as an independent reference. Specifically, 1 H1 WT and 1 HUES8 WT hPSC clone were each infected separately with 12 different LARRY barcodes, sorted, and cryopreserved. WT cells were then differentiated and collected at the following time points: day −1, 3, 5, 7, 9, 10, 11, 12, 14, 16, 18, and 26. Cells were cryopreserved in Bambanker cell freezing medium after collection.

On the day of scRNA-seq library preparation, frozen cells were thawed, sorted, and multiplexed such that each 10x Genomics reaction contained WT cells from multiple time points. Barcode sequences enabled discrimination of cells from different time points and cell backgrounds. cDNA and targeted libraries were generated and sequenced as described above.

#### Flow cytometry

Cells were dissociated using TrypLE Select, washed in FACS buffer (5% FBS and 5 mM EDTA in PBS) and then incubated with Live-Dead Fixable Violet Dead cell stain (Invitrogen, L34955, 1:1,000 dilution) in FACS buffer for 15 minutes at room temperature. Afterwards, cells were fixed with Fixation/Permeabilization reagent (Invitrogen, 00522356/00512343) for 1 hour at room temperature. Intracellular protein staining was performed by sequential incubation with primary and secondary antibodies, each for 1 hour in Permeabilization Buffer (Thermo Fisher Scientific, 00833356). After staining, cells were resuspended in FACS buffer for analysis using BD LSRFortessa. Data analysis and figures were generated using FlowJo v.10. The antibodies are listed in Supplementary Table 11.

#### Microarray of *Rfx6* knockout mouse pancreata

Animal experiments were supervised by G. Gradwohl (agreement N°C67-59 approved by the Direction des Services Vétérinaires, Strasbourg, France) in compliance with the European legislation on care and use of laboratory animals. The generation and maintenance of mouse strains has been described previously [77]. Two genetic comparisons were included in the microarray analysis: (1) *Rfx6fl/fl; Ngn3-Cre* conditional knockouts with *Rfx6fl/+* littermates as controls and (2) *Rfx6-/-* full knockouts with WT as controls.

Embryos were dissected at E15.5 for the pancreas. Individual pancreata were harvested and dissociated in 1 ml of Tri Reagent (Sigma) using a 26G needle. Samples were snap frozen in liquid nitrogen and stored at −80°C until genotyping. RNA extraction was then performed using RNAeasy Minikit (Qiagen). RNA from 3-4 biological samples per condition was hybridized on Agilent microarrays (Gene Expression Microarray, 8×60K: SurePrint G3 Mouse GE 8×60K G4852A) by the Biopuce platform of the IGBMC. Microarray analysis was performed as described previously [78]. The processed expression data are provided in Supplementary Table 9.

#### Generation of *ISL1-/-* hPSC clonal lines

*ISL1-/-* clones were generated in H1 hPSC background using a knockin-knockout strategy. Cells were co-transfected with Cas9-ISL1 gRNA plasmid and 2 donor plasmids designed to insert PGK-neomycin and PGK-puromycin resistance cassettes in the *ISL1* exon 3 using Lonza 4D nucleofector with P3 Primary Cell 4D-Nucleofector X Kit S (Lonza, V4XP-3032) [79]. 3 days after transfection, cells were treated with 200 μg/ml neomycin and 1 μg/ml puromycin concurrently for 3 days to select for biallelic knockin cells. Single-cell clones were then picked and evaluated with PCR. Clones that were positive for both the puromycin and neomycin junction PCRs, showed the expected knock-in product in the across PCR, and lacked the WT allele were selected as biallelic knock-in clones. The gRNA target sequence and genotyping primers were as follows. *ISL1* gRNA: acactcgatgtgataca. PCR-F1: TCCCCTTCTCCTACCTCCTT. PCR-R1: GGGGAATCAAGGGAGACAGT. Puro-R: gtgggcttgtactcggtcat, Neo-R: ctcgtcctgcagttcattca.

#### *ISL1* overexpression experiment

EF1a-ISL1-IRES-BFP lentiviral overexpression plasmid was generated by cloning the *ISL1* open reading frame (Addgene #142382) and IRES-BFP (Addgene #85449) into the LARRY backbone (Addgene #140024). An EF1a-BFP construct was generated in parallel as a negative control. Plasmids were transfected into 293T cells to produce lentivirus, which was subsequently concentrated to 100× by ultracentrifugation at 25,000 rpm 4°C for 90 minutes.

On day 11 of differentiation, cells were dissociated using TrypLE, and approximately 6 × 10⁶ cells were resuspended in 5 ml S5 medium supplemented with 100 μl ISL1-BFP or BFP concentrated lentivirus, 10 μM Y-27632, and 6 μg/ml protamine sulfate. The cell suspension was transferred to a single well of an ultra–low attachment 6-well plate and incubated on an orbital shaker at 100 rpm. On day 18 of differentiation, BFP+ cells were sorted by flow cytometry. RNA was extracted using the Quick-RNA Purification Kit (Zymo Research, R1055) and submitted for SMARTer bulk RNA-seq (SMART-Seq v4 + KAPA EvoPrep) through the Integrated Genomics Operation Core at MSKCC.

### II. Data analysis

#### Pre-processing of the 10x scRNA-seq data

Transcriptome and barcode libraries were processed using Cell Ranger (v.7.1.0) with default parameters to generate gene-cell count matrices. Downstream analyses were performed in Seurat (v.5.0.3). To ensure high-quality single-cell profiles, we applied multiple filtering criteria. Cells were excluded if they contained fewer than 10 LARRY barcode UMIs per cell, exhibited multiple barcodes with secondary barcode counts at least 50% of the primary barcode, expressed fewer than 200 or more than 7,000 genes, or had more than 10% of total reads derived from mitochondrial genes. In addition, 1 cluster that was enriched for cells with high mitochondrial content after dimension reduction was also removed. Following these quality control steps, approximately 40-60% of cells were retained, yielding 87,173 cells from KO pool and 74,606 cells from external WT dataset. These retained cells were subsequently assigned genotypes based on barcode information for downstream analysis.

#### scRNA-seq normalization, integration, and cell type annotation

For the transcriptome libraries derived from external WT cells, ambient RNA contamination was observed, likely introduced during the pooling of cells from multiple differentiation time points prior to scRNA-seq. This contamination resulted in low-level background expression of SC-islet genes (e.g, *CHGA*, *INS*) in early-stage cells. To address this, doublets were first identified and removed using DoubletFinder (v2.0.4), and ambient RNA contamination was subsequently corrected using DecontX implemented in the celda R package (v1.18.1). Cells with high residual contamination were further removed based on their DecontX contamination scores. To avoid over-filtering, the contamination score threshold was adjusted such that the proportion of INS+ cells remained below 3%, matching the ∼3% INS+ population observed in day 11 internal WT cells.

The knockout village dataset and cleaned external WT dataset were then integrated using Seurat’s reciprocal PCA (RPCA) method with default parameters. Normalization, scaling, principal component analysis (PCA), clustering, and UMAP visualization were performed on the top 2,000 highly variable genes and top 30 principal components. Because the rate of cell proliferation naturally decreases during differentiation, cell cycle effects were not regressed out to preserve this biologically meaningful variation. Cell type identities were finally assigned to clusters based on the top differentially expressed genes and established cell type marker genes described in the literature.

#### scRNA-seq pseudo-bulk PCA

Pseudo-bulk expression profiles were generated by aggregating raw counts from Seurat RNA assay across cells of each clone x time condition. Sample-level metadata was constructed for DESeq2 (v1.34) analysis using the design formula ∼ genotype + time. PCA was performed on the top 1,000 most variable genes, and the first two principal components were visualized with ggplot2.

#### Gene expression variance analysis

To quantify the variance composition, pseudo-bulk samples were created for each clone x time condition that had at least 30 cells. Genotypes with at least 2 clones passing the cell number criterium were included in the analysis. For each time point, we considered top 5,000 highly variable genes identified using the ‘FindVariableFeatures’ in Seurat to capture the variance composition pattern. For each selected highly variable genes, we fitted a series of nested linear models using a group of predictors in the following sequential order: library size as quantified by the total count per pseudo-bulk sample, genetic background, and genotype. Variance explained by each factor was quantified by adjusted R2 from the sequence of linear models.

#### Cell type composition analysis

Based on differentiation trajectories, we grouped cell types into three categories: 1. intended cell types: cells belonging to the desired islet lineage and captured at the expected time point (e.g., hPSC at D-1, DE at D3, PFG at D7, PP at D11, and SC-islet cells (EnP, SC-β, SC-α, SC-δ, and SC-EC) at D18). 2. delayed cell types: cells from the desired islet lineage but captured at a later-than-expected time point (e.g., ESC and DE cells at D7). 3. alternative cell types: cells from alternative lineages (e.g., stromal, endothelial, liver, or PGP cells) captured at any time point.

To quantify mutant effects on cell type compositions, we first calculated the proportion of each cell type/category per time point for each clone. Clones with ≤25 cells at a given time point were excluded. For each cell type/category and time point, we then fitted a linear regression model: log(cell type proportion) ∼ background + genotype. The *cell type proportion* was transformed by DR_data() before log to avoid near zero values. The *background* represents genetic background (H1 or HUES8). Adjusted p < 0.05 was considered statistically significant.

#### β cell state score

To define β cell state, we first identified β cell marker genes by performing differential gene expression analysis using FindMarkers() function from Seurat. This compared WT SC-β cells to all other WT cell types on day 18, resulting in 95 β cell marker genes (adjusted p < 0.05 and abs(log₂(fold change)) > 1). For β cell state scoring, all SC-β cells and a randomly sampled 20% of other WT cells from day 18 (serving as negative controls) were selected. The scmageck_eff_estimate() function from scMAGeCK (v1.9.2) was then used to calculate a β cell state score for each cell based on expression of the 95 marker genes, using default parameters. Each cell received a score ranging from 0 to 1, with lower scores indicating stronger transcriptional deviation from WT SC-β cells.

#### Differential gene expression (DEG) analysis of mutant genotypes vs WT

DEGs were identified by comparing each mutant genotype to WT using FindMarkers() function, with the cutoff of adjusted_p < 0.005 and abs(avg_log2FC) > 0.5. GO enrichment analysis was performed using Metascape[4] on upregulated and downregulated genes. For *NEUROD1-/-* SC-β, we further compared the DEGs against published bulk RNA-seq. Upregulated and downregulated gene sets were generated from Neurod1-/- E17-sorted pancreatic endocrine cells using p_adj < 0.05 and abs(log2FC) > 0.7 [30] and from E15.5-sorted endocrine cells using p_adj < 0.05 and abs(log2FC) > 0.5 [32]. Gene Set Enrichment Analysis (GSEA) was performed using GSEA v4.3.3 with the pre-ranked option based on log2FC values.

#### Curation of T2D risk genes and enrichment analysis

To generate genes credibly associated with Type 2 Diabetes (T2D), we utilized publicly available data from the Open Targets Platform (https://www.opentargets.org/, March 2024 release). The Experimental Factor Ontology (EFO) term corresponding to T2D (MONDO_0005148) was used to query the Open Targets Genetics portal and retrieve all related Genome-Wide Association Studies (GWAS). Unique study identifiers from the query were then used to download comprehensive datasets for each study. For each genetic locus, we extracted the credible set of variants, defined as the set of variants with a 95% probability of containing the causal variant driving the association signal. Variants were linked to putative causal genes using the Variant-to-Gene (V2G) scores provided by Open Targets. These scores are derived from the Locus-to-Gene (L2G) model, which integrates functional genomics, molecular QTLs, and chromatin interaction data to estimate the likelihood (ranging from 0 to 1) that a given gene is causally responsible for the GWAS signal. Datasets containing study metadata, credible set variants, and V2G scores were integrated and deduplicated to produce a non-redundant list of T2D-associated genes and variants. A total of 485 high-confidence T2D risk genes (L2G score > 0.5) that were also detected in our scRNA-seq dataset were retained for downstream enrichment analysis.

To assess whether T2D risk genes were enriched in DEGs of any diabetes mutant genotype, we performed an odds ratio-based enrichment analysis. For each DEG list, a 2×2 contingency table was constructed comparing overlap with the T2D gene set:

A = number of genes in the DEG list overlapping with the T2D risk gene set
B = number of genes in the DEG list not overlapping with the T2D risk gene set
C = number of T2D risk genes not overlapping with the DEG list
D = number of non-T2D risk genes outside of the DEG list

The odds ratio (OR) was calculated as (A/B) / (C/D), with the standard error (SE) estimated as 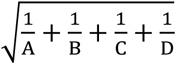, and the absolute z-score (z_abs) as |log(OR)/SE|. The corresponding p-value was computed by scipy.stats.norm.sf(Z_abs), and all p-values were adjusted for multiple testing using the Benjamini–Hochberg false discovery rate method.

#### Extracting gene programs by non-negative matrix factorization

We used non-negative matrix factorization (NMF) to identify patterns of gene co-expression across the single-cell dataset. Starting from the filtered and normalized expression values, we filtered for the top 5,000 most highly variable genes using the sc.pp.highly_variable_genes function in Scanpy (v1.10.3) and formed a filtered and normalized expression matrix *X* of dimension 161,779 x 5,000. Writing *N*, *G*, *k* to be the number of cells, genes, and the NMF approximation rank respectively, we used sk.decomposition.NMF function to find two factor matrices *U* (cell-by-program, *N* × *k*) and *V* (gene-by-program, *G* × *k*) such that *X* ≈ *UV*^*T*^. The choice of the number of programs or components, *k*, is detailed below. We used the NMF implementation of scikit-learn [80] and used a coordinate descent algorithm and the Nonnegative Double Singular Value Decomposition (NNDSVD) initialization [81], which is a structured, non-negative and deterministic initialization algorithm for NMF.

Once activities *U* and programs *V* were fit, each of the 5,000 highly variable genes were scored against gene programs. For each gene, expression values were first z-scored and regressed against the columns of *U*. This produced coefficients for each pair of program and gene. For a given program, the genes with the most positive (resp. most negative) coefficients correspond to the most correlated (resp. anticorrelated) genes.

#### Choosing the number of components

To choose the number of components (*k*), we evaluated a range of values from 4 to 72 (i.e., *k* = 4, 8, 12, …, 72). We selected *k* = 36 for all downstream analyses based on two criteria: (1) gene coverage, measured by aggregating the top 300 marker genes from each gene program; (2) functional diversity, assessed by the total number of unique Gene Ontology (GO) terms enriched among top 100 marker genes of each gene program. GO enrichment was performed by enrichGO() function from package clusterProfiler (v4.12.6). Although *k* = 36 didn’t yield the maximum number of gene or GO term counts, it captured the majority of functional diversity in the dataset. As higher *k* values would increase redundancy across programs, *k* = 36 represented a balanced choice between resolution and interpretability.

#### Gene program annotations

To facilitate biological interpretations, we manually curated and grouped gene programs into 12 functional categories based on enriched GO terms. GO enrichment analysis was performed on top 100 marker genes of each gene program ranked by the z-score regression coefficients, using clusterProfiler and Metascape [82]. We empirically found that top 100 genes provided clearer functional characterization of gene programs compared to larger sets (e.g., top 200 or 300). To quantitatively assess similarity among gene programs, we calculated pairwise Jaccard similarity based on their top 100 genes. This provided an unbiased measure of relatedness across functional categories. The grouped gene programs were visualized using Cytoscape v3.10.3.

#### Assessing gene program activities in human fetal scRNA-seq

To evaluate the relevance of identified gene programs in human fetal tissues, we analyzed a published human fetal scRNA-seq dataset [39]. Cells from gestational weeks 11-13 were selected and then filtered to retain only the major representative cell types for each tissue, as defined by their original cell type annotations from CZ CELL X GENE data portal (https://cellxgene.cziscience.com/collections/38833785-fac5-48fd-944a-0f62a4c23ed1). This process resulted in 16 representative cell types from 15 tissues. Gene program activities were quantified using the top 100 genes from each program with the AddModuleScore_UCell() function from UCell (v2.8.0), which generated single-cell-level activity scores. Mean activity scores were then calculated per cell type and scaled to obtain the relative activity of each gene program across fetal cell types.

#### Gene regulatory network inference by SCENIC+

We applied SCENIC+ to infer gene regulatory network among endocrine cells, including SC-α, SC-β, SC-δ, SC-EC, and PP cells. The sc-multiome data were processed using the pycisTopic Python package. Specifically, pseudo-bulk samples were generated for each cell type, followed by consensus peak calling using MACS2. A topic model was fitted to identify topic-specific regions, which were then aggregated with cell-type-specific differentially accessible regions for motif enrichment analysis using pycisTarget. The SCENIC+ pipeline was subsequently applied using default parameters.

#### Inference of transcription factors for each gene program

The SCENIC+ predicted in total 139 TF regulons, comprising 7473 direct target genes. Among these TF regulons, 45 contained more than 100 target genes and were defined as core TFs in endocrine regulation. To assess relationships among core TF regulons, we quantified the overlap between their target gene sets using Fisher’s exact test, and visualized significant associations (odds ratio > 1, p_adjusted < 0.001) as a network by Cytoscape v3.10.3. Next, to identify potential upstream regulators of each gene program, we performed Fisher’s exact test to evaluate the enrichment between the top 100 genes of each gene program and the core TF regulons. TFs showing significant enrichment (p_adjusted < 0.001) were inferred as candidate upstream regulators. All p-values were adjusted for multiple testing using the Benjamini–Hochberg false discovery rate correction.

#### Linear regression analysis

EnP cells were extracted from integrated Seurat object using subset() function. A total of 6422 genes with non-zero counts in at least 20% EnP cells were selected for linear regression analysis. Genotypes with 10 or fewer cells were excluded. To assess how gene expression in EnP correlate with SC-β/SC-EC cell fate bias, we performed gene-wise linear regression across mutant genotypes. For each gene, the model was defined as: 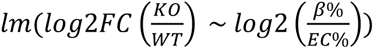, where each data point corresponds to a mutant genotype.

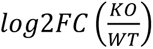 represents the log2 fold change in EnP gene expression between mutants and WT, and 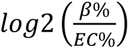 represents the SC-β/SC-EC cell fate bias in mutants on day 18. The regression coefficient and p value of individual genes were saved in Supplementary Table 8.

#### Bulk RNA-seq analysis

RNA-seq reads were aligned to the human reference genome (GRCh38, primary assembly) using STAR (v2.7.11b) with the aid of its genome indexing function and gene annotation from GENCODE (v44). Following alignment, a gene-level read count matrix was created using featureCounts (v2.1.1) and the same gene annotation. Normalization, PCA and DEGs were generated by DESeq2 (v1.34.0). Analyzed RNA-seq data (normalized counts) are presented in Supplementary Table 10. DEGs were identified based on cut-off abs(log2(FC)) > 1 and p_adjusted < 0.05. GO analysis was performed using Metascape [82] and GSEA was performed using clusterProfiler.

## Data availability

The knockout village scRNA-seq raw sequencing data are available from the ENA under accession ERP168459, and the processed read counts are available at the GEO under accession GSE313516. The processed Seurat object and metadata, including cell type and genotype annotations, are available on Zendo (https://doi.org/10.5281/zenodo.17824935). ISL1 overexpression bulk RNA-seq data are available at GSE313800 (reviewer token: ufwlukewlxalzst). Previously published human fetal scRNA-seq data and SC-islet sc-multiome data that were re-analyzed here are available at GSE134355 and GSE199636, respectively.

## Acknowledgments

We acknowledge the assistance from the following Memorial Sloan Kettering Cancer Center (MSKCC) Cores: Antibody & Bioresource, Flow Cytometry, Gene Editing & Screening, Integrated Genomics Operation, and Stem Cell Research. We thank R. Garippa, S. Mehta and Zakheim for assistance with LARRY barcode library expansion, L. Hung, K. Y. Yeung, A. Shivalikanjli, and H. Cho for assistance with scRNA-seq data preprocessing and data brokering and deposition as part of the MorPhiC consortium effort, L. Studer, A.-K. Hadjantonakis, J. Choi and members of D.H.’s laboratory for insightful discussions. We acknowledge the following funding resources: D.H. is supported by grants from NIH (UM1HG012654, R01HD111256, and R01DK096239), MSKCC Cancer Center Support Grant from NIH (P30CA008748). W.L. is supported by grants from NIH (R01HG010793, R01HL168174), the National Cancer Institute - Cancer Center Support Grant (CCSG) - P30CA134274, startup funding support from UM-IHC, Montgomery County, Maryland, and the University of Maryland Strategic Partnership: MPowering the State, a formal collaboration between the University of Maryland, College Park and the University of Maryland, Baltimore. G.G. is supported by grants from the Agence Nationale pour la Recherche (ANR Rfx-PancInt) and the Fondation pour la Recherche Médicale (FRM). D.L. was supported by a Beatrice P. K. Palestin Fellowship and a Bruce Charles Forbes Fellowship. D.Y. was supported by a postdoctoral fellowship from a NYSTEM training grant (DOH01-TRAIN3-2015-2016-00006). G.D. was supported by an NIH T32 Training Grant and a Frank Lappin Horsfall Jr Fellowship. A.I. is supported by a Medical Scientist Training Program grant from the NIGMS of the National Institutes of Health under award number T32GM152349 to the Weill Cornell/Rockefeller/Sloan Kettering Tri-Institutional MD-PhD Program.

## Author contributions

D.L., D.H., W.L., W.S., K.D., and D.Y. designed experiments and interpreted results. D.L. performed most experiments and analyzed the results. D.Y. assisted with village method establishment. W.L. supervised and B.S. performed scRNA-seq preprocessing and integration. W.S. supervised and Z.L. modeled gene expression variance, cell type composition change and islet TF regulomes. X.Q. supervised and S.Z. assisted with gene program analysis. K.D. supervised and T.F. assisted with T2D-related enrichment analysis. G.G. supervised and J.P. performed microarray on mouse fetal pancreata. T.Z. supervised and A.Z. generated ISL1-/- hPSC lines. D.Y., G.D., and C.S. assisted with the generation of hPSC knockout lines. R.L. assisted with scRNA-seq experiment. J.Z., A.I., N.H., and B.O. assisted with additional experiments. V.M. assisted with gene regulatory network analysis. D.L. and D.H. wrote the manuscript. All authors provided editorial advice.

## Competing interests

Authors declare that they have no competing interests.

---

**Supplementary Fig. 1.**
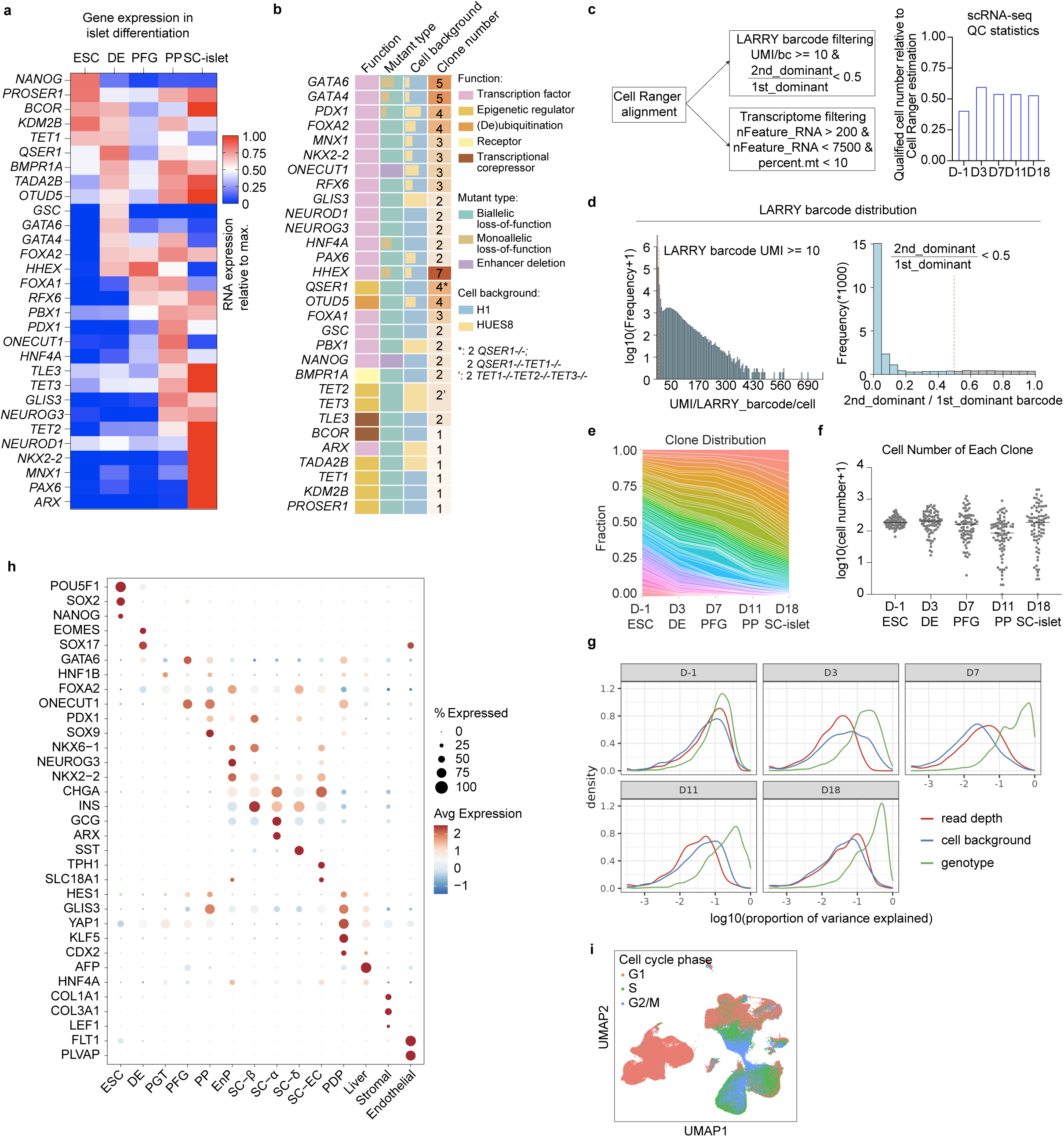
Statistics and quality control related to the knockout village. **a.** The expression dynamics of perturbed genes during hPSC-derived islet differentiation, based on in-house bulk RNA-seq. **b.** The list of perturbed genes, categorized by the function, mutant type, cell background and clone number in the knockout village. **c.** Left: quality control strategy on LARRY barcode libraries and transcriptome libraries. Right: The fraction of cells passing both filtering steps. **d.** Histogram showing the distribution of LARRY barcode read counts before filtering steps. **e.** Stacked line plot showing the clone representation during differentiation. Each line represents a clone. **f.** Cell numbers of individual clones during differentiation. The lines represent the median cell number. **g.** Histograms showing the contribution of read depth, cell background, and genotype to gene expression variance. **h.** Dot plot showing the expression of marker genes associated with each cell type. **i.** UMAP embedding of scRNA-seq dataset, annotated by cell cycle phase.

**Supplementary Fig. 2.**
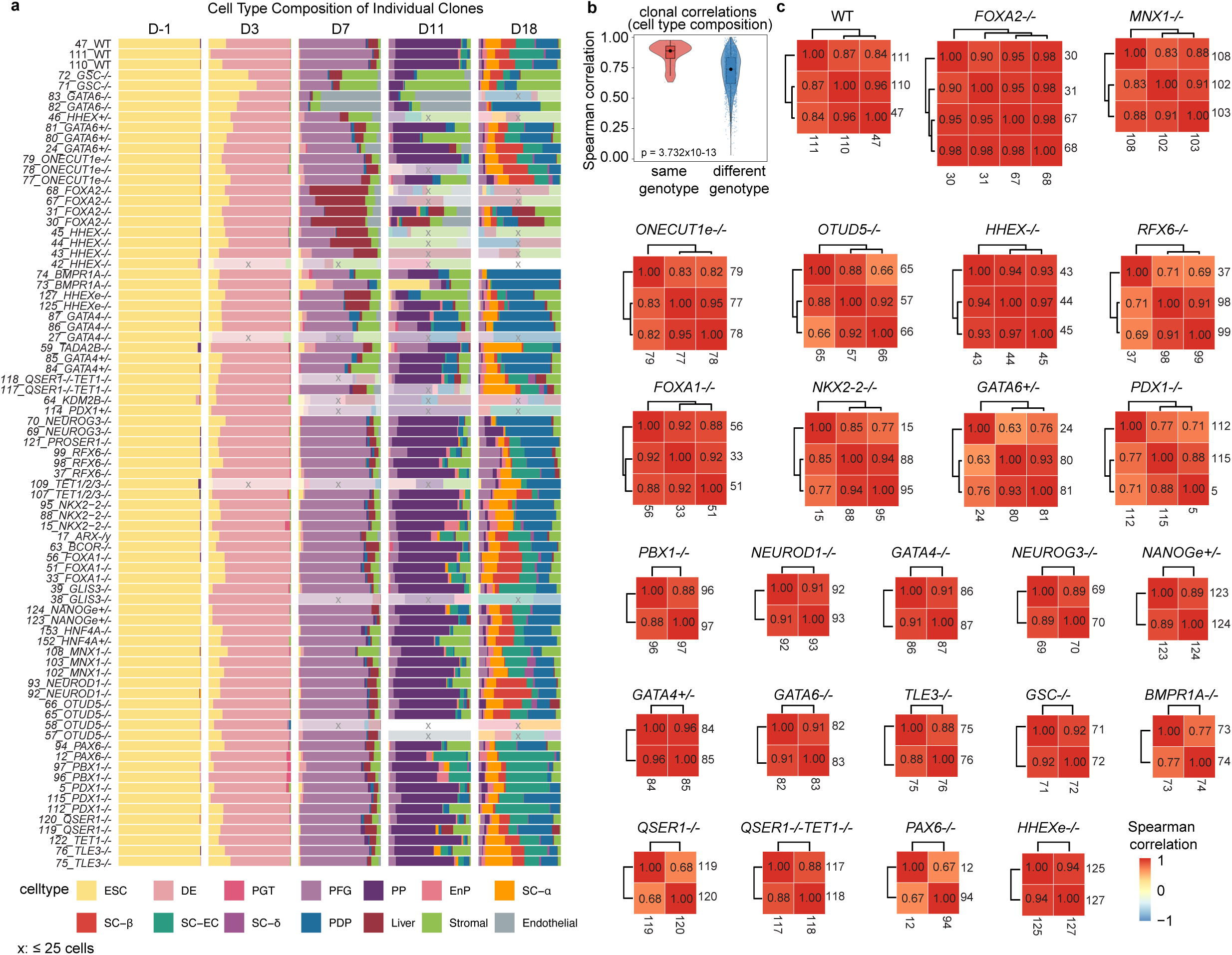
Clones from the same genotype exhibit comparable cell type compositions. **a.** Cell type compositions of individual mutant clones during hPSC-derived islet differentiation. Clones with 25 or fewer cells were marked with ‘x’ (not analyzed). **b.** Violin plot showing the distribution of Spearman correlation coefficients between clone pairs from the same or different genotypes, based on their cell type compositions. Each dot represents a clone pair. Box plots indicate the median and interquartile range (25th–75th percentile). **c.** Heatmap showing the Spearman correlation coefficients between clone pairs from the same multi-clone genotypes.

**Supplementary Fig. 3.**
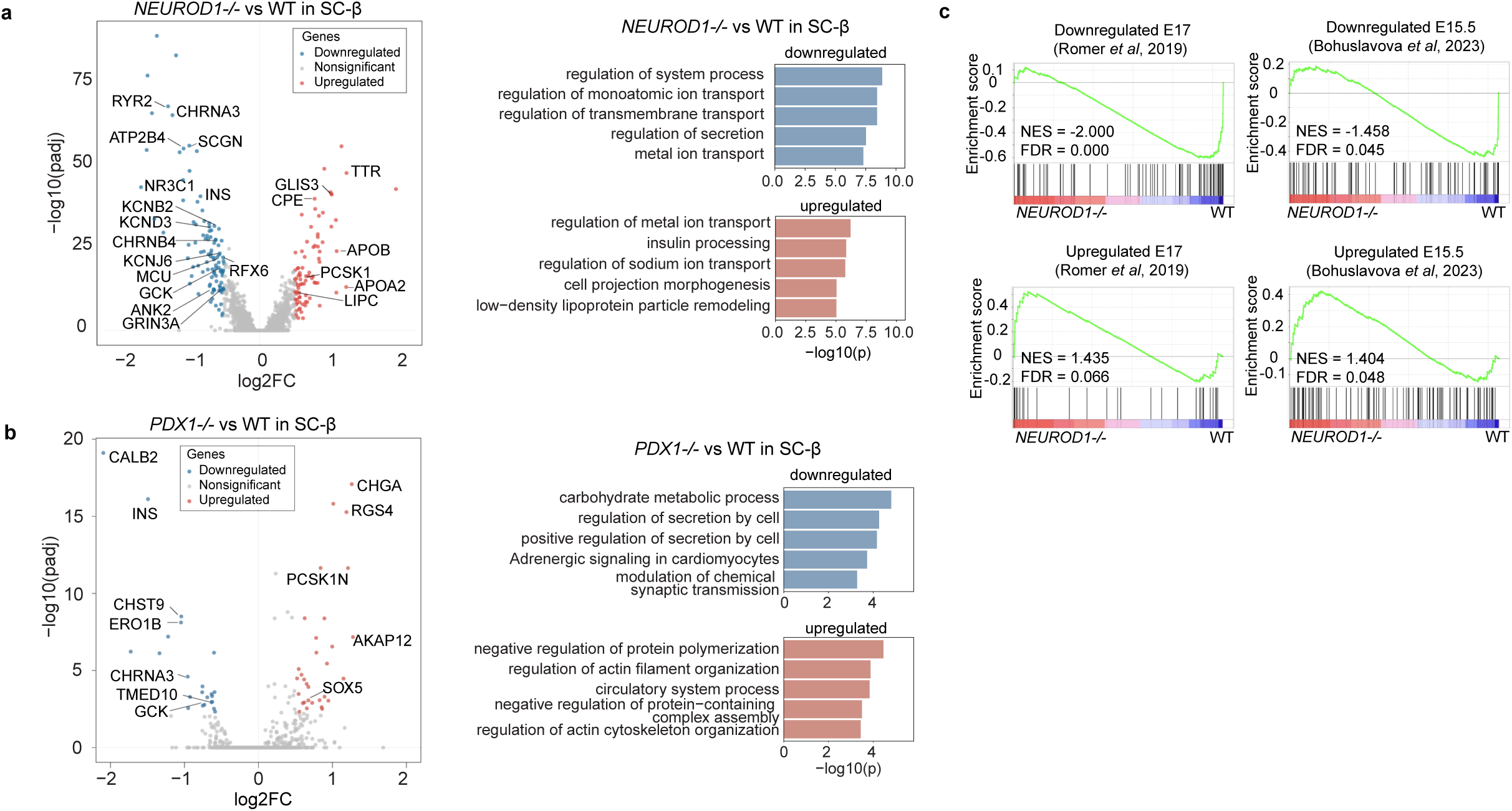
Loss of *PDX1* or *NEUROD1* alters SC-β state. **a-b.** Left: volcano plots showing differential gene expression in SC-β cells for *PDX1-/-* vs WT (a) and *NEUROD1-/-* vs WT (b). Significance thresholds: abs(log2(FC)) > 0.5 and *p*_adjusted < 0.005. Right: top 5 enriched gene ontology terms in down- and upregulated genes from *PDX1-/-* vs WT (a) and *NEUROD1-/-* vs WT (b). **c.** GSEA of *NEUROD1-/-* vs WT SC-β cells, with genes ranked by log2 fold change and tested for enrichment of down- and upregulated gene sets derived from mouse fetal endocrine cells (*Neurod1-/-* vs WT).

**Supplementary Fig. 4.**
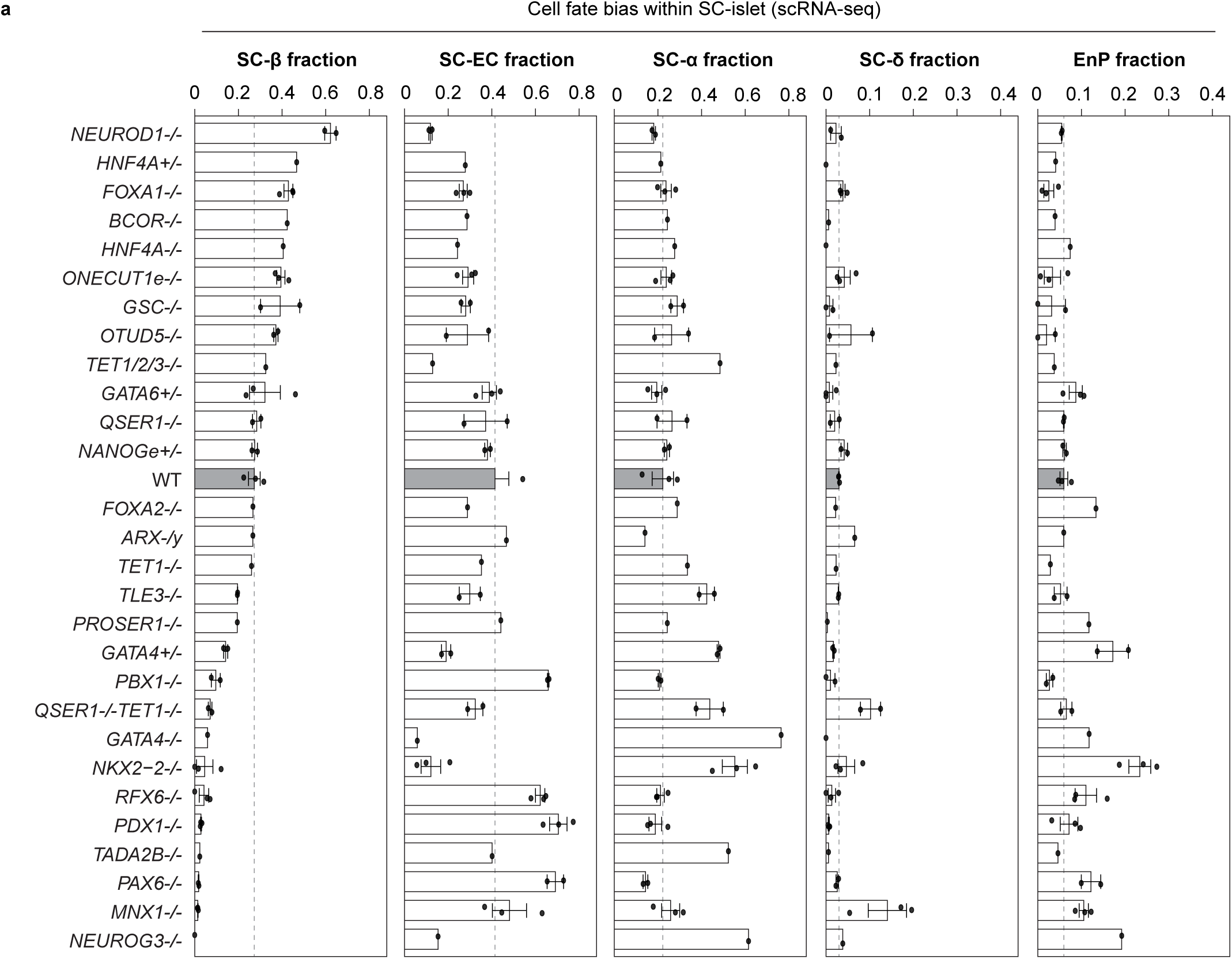
Effects of mutant genotypes on cell type compositions within endocrine population. **a.** Fractions of individual endocrine cell types within total endocrine cells on day 18. Each dot represents a clone with over 25 endocrine cells on day 18. Only genotypes with at least one clone passing the cell number filter are shown. Dash lines indicate the mean fraction across WT clones.

**Supplementary Fig. 5.**
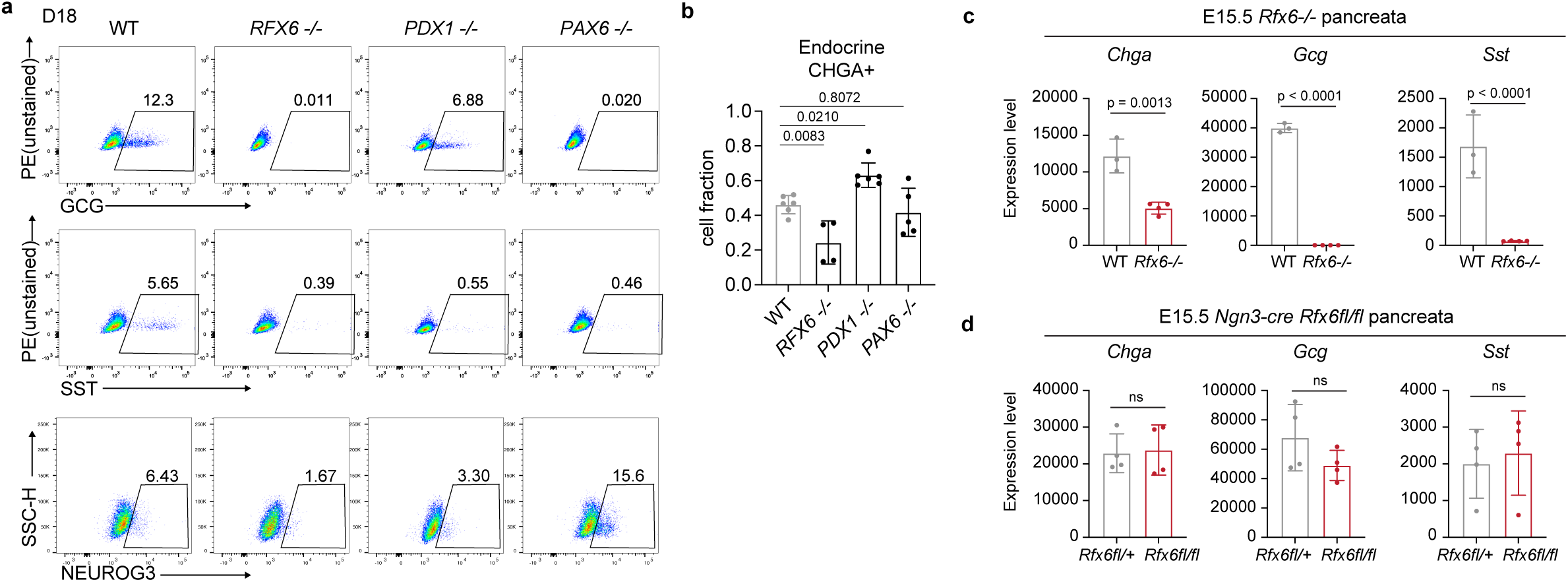
Effects of *RFX6-/-*, *PDX1-/-*, and *PAX6-/-* on endocrine cell fates. **a.** Representative flow cytometry plots of SC-α (GCG), SC-δ (SST) and EnP (NEUROG3) markers in WT, *RFX6-/-*, *PDX1-/-*, and *PAX6-/-* cells on day 18. **b.** Quantification of CHGA+ fraction on day 18 from flow cytometry analysis. n = 6 (WT and PDX1-/-), n=5 (PAX6-/-) and n=4 (RFX6-/-) independent differentiations. *P* values were calculated using one-way ANOVA test with Dunnett’s multiple comparisons test. **c-d.** mRNA level of pan-endocrine (*Chga*), α cell (*Gcg*) and δ cell (*Sst*) markers in E15.5 WT and *Rfx6-/-* pancreata (c), and in E15.5 *Ngn3-cre Rfx6fl/+* and *Rfx6fl/fl* pancreata (d), measured by microarray. n = 4 (*Rfx6-/-, Ngn3-cre Rfx6fl/+,* and *Ngn3-cre Rfx6fl/fl*), n=3 (WT) samples.

**Supplementary Fig. 6.**
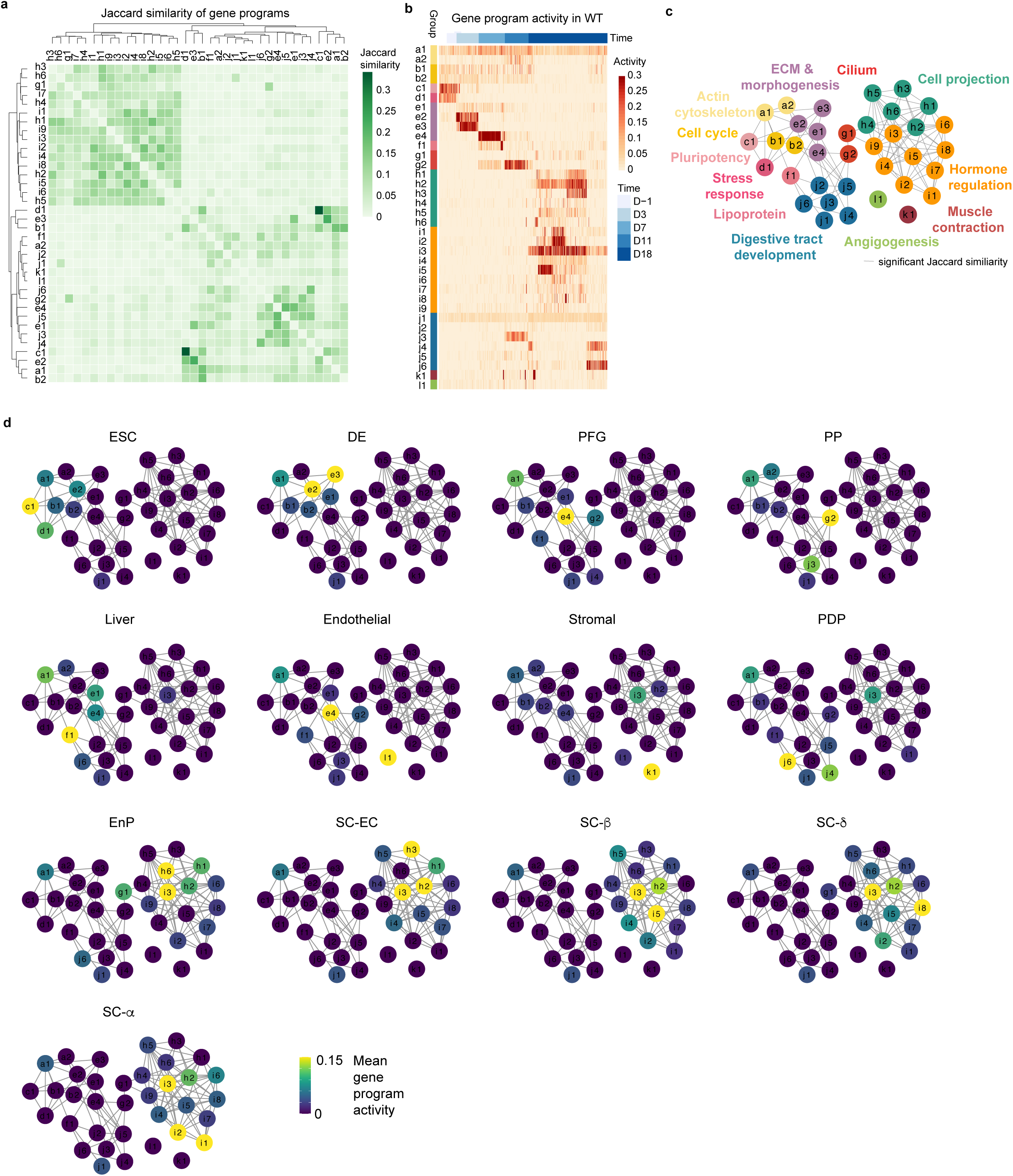
Gene programs uncover biological processes associated with differentiating cell types. **a.** Jaccard similarity between gene program pairs, calculated using the top 200 genes from each program. **b.** Gene program activity across WT cells. Each column represents a cell, ordered by differentiation time points. To facilitate interpretations, gene programs were manually grouped into 12 categories based on gene ontology analysis. **c.** Network view of gene programs colored by gene ontology-based categories. Nodes represent gene programs. Edges connect programs with Jaccard similarity above the null distribution. **d.** Network view of mean gene program activity in WT cell types.

**Supplementary Fig. 7.**
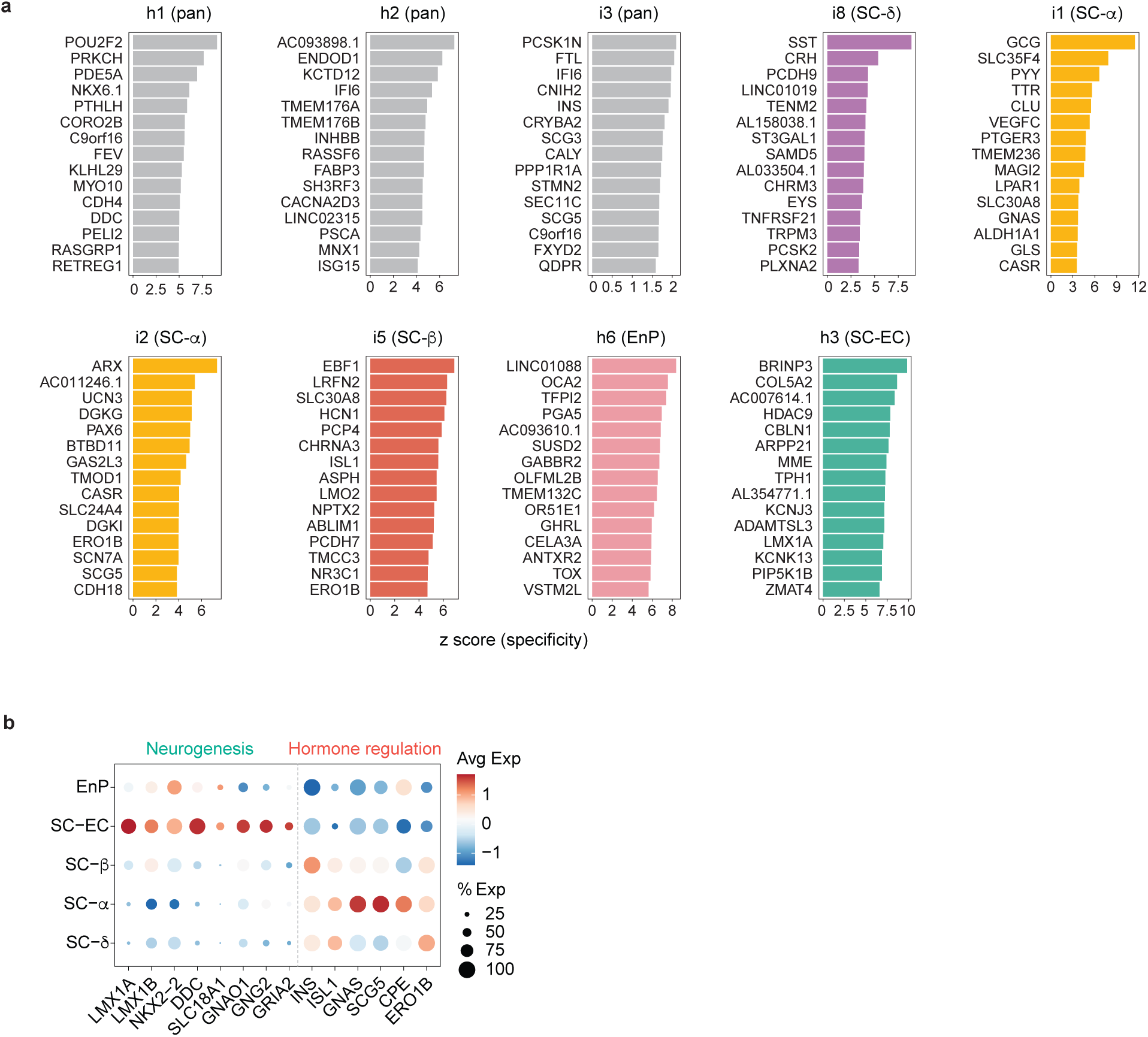
Gene programs reveal both shared and cell type-specific signatures in the endocrine lineage. **a.** Top 15 genes of endocrine-associated gene programs, ranked by z-score specificity. **b.** Expression of representative neurogenesis and hormone regulation markers in WT endocrine cell types.

**Supplementary Fig. 8.**
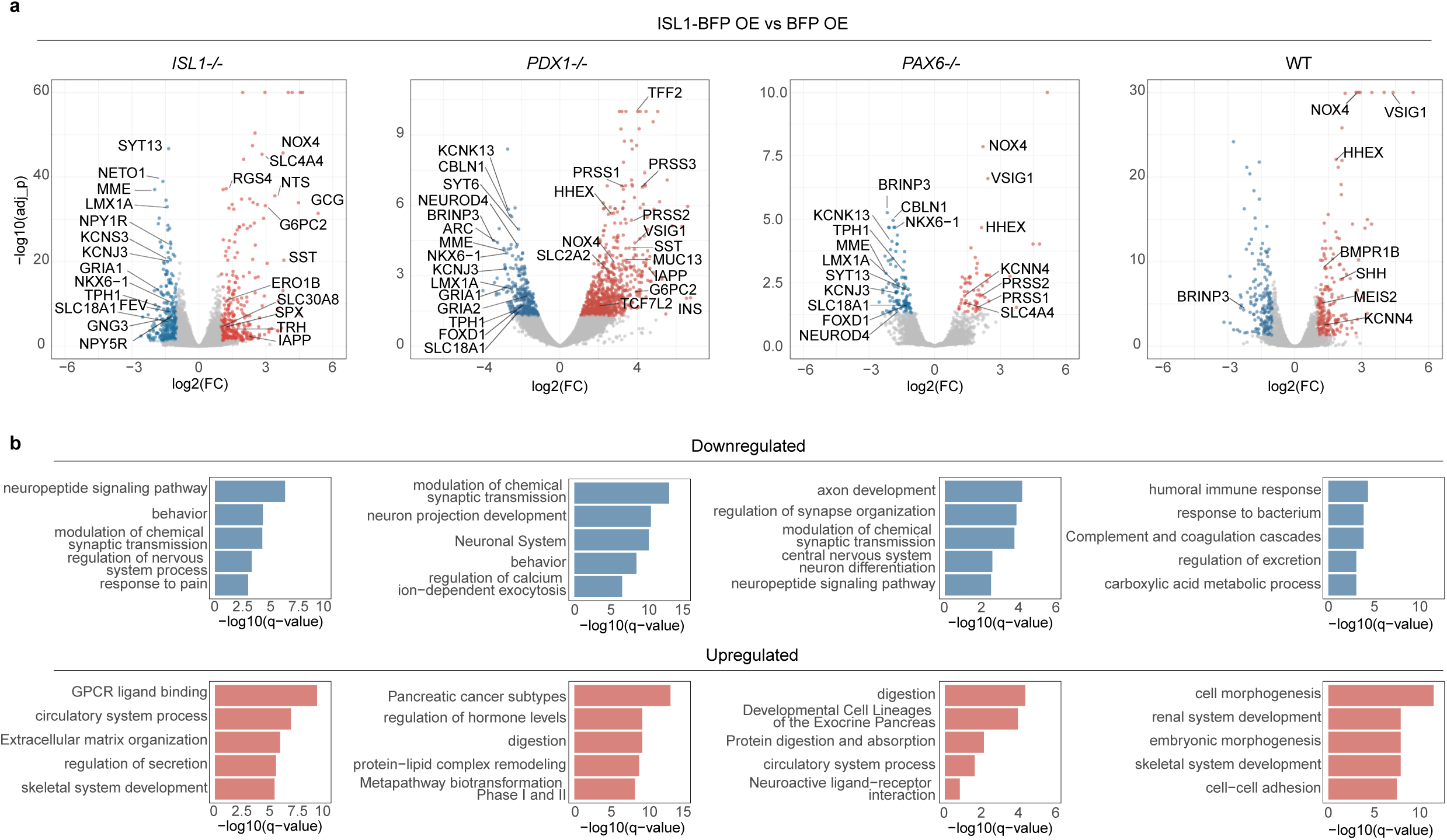
*ISL1* overexpression reduces neuron-related gene expression during islet differentiation. **a.** Volcano plots showing differential gene expression in *ISL1-BFP* OE vs *BFP* OE (control) across genotypes. Significance thresholds: abs(log2(FC)) > 1 and p_adjusted < 0.05. **b.** Gene ontology analysis of top five enriched terms among upregulated and downregulated genes in *ISL1-BFP* OE vs *BFP* OE (control) across genotypes.

## Supplementary Tables

**Table 1.** Genotypes and corresponding genotyping primers for hPSC lines in the knockout village.

**Table 2.** Cell type composition of individual lines in knockout village.

**Table 3.** The list of 95 genes for SC-β cell state score analysis.

**Table 4.** The list of T2D risk genes curated from Open Targets database.

**Table 5.** Specificity z-scores for genes in each gene program.

**Table 6.** Top 10 gene ontology terms enriched in each gene program.

**Table 7.** SC-islet TF regulons inferred by SCENIC+.

**Table 8.** Linear regression coefficients between EnP gene expression and SC-EC/SC-β cell fractions on day 18.

**Table 9.** mRNA counts from fetal pancreata of WT vs *Rfx6-/-* and *Ngn3-cre Rfx6fl/+* vs *Ngn3-cre Rfx6fl/fl* on day18, based on microarray data.

**Table 10.** Normalized read counts from bulk RNA-seq of ISL1 overexpression in WT, ISL1-/-, PDX1-/-, and PAX6-/- on day 18.

**Table 11.** Antibody list.

## Notes

### Competing Interest Statement

The authors have declared no competing interest.

